# Computational design of dynamic biosensors for emerging synthetic opioids

**DOI:** 10.1101/2025.05.15.654300

**Authors:** Alison C. Leonard, Chase Lenert-Mondou, Rachel Chayer, Samuel Swift, Zachary T. Baumer, Ryan Delaney, Anika J. Friedman, Nicholas R. Robertson, Norman Seder, Jordan Wells, Lindsey M. Whitmore, Sean R. Cutler, Michael R. Shirts, Ian Wheeldon, Timothy A. Whitehead

## Abstract

Nitazenes are an emergent class of synthetic opioids that rival or exceed fentanyl in their potency. These compounds have been detected internationally in illicit drugs and are the cause of increasing numbers of hospitalizations and overdoses. New analogs are consistently released, making detection challenging – new ways of testing a wide range of nitazenes and their metabolic products are urgently needed. Here, we developed a computational protocol to redesign the plant abscisic acid receptor PYR1 to bind diverse nitazenes and maintain its dynamic transduction mechanism. The best design has a low nanomolar limit of detection *in vitro* against nitazene and menitazene. Deep mutational scanning yielded sensors able to recognize a range of clinically relevant nitazenes and the common metabolic byproduct in complex biological matrices with limited cross-specificity against unrelated opioids. Application of protein design tools on privileged receptors like PYR1 may yield general sensors for a wide range of applications *in vitro* and *in vivo*.

## Introduction

The rise in overdose deaths from synthetic opioids has been well-documented in the US and worldwide, with prevalence rising in the 2010s to account for well over half of documented overdose deaths in the US in 2022 (*1–3*). Synthetic opioids such as fentanyl are inexpensive to produce and highly potent, with fentanyl 50 to 100 times more potent than heroin and morphine (*2*, *4*), and thus are often mixed with other drugs to cheaply increase strength and addictivity (*2*, *3*). An emerging class of novel synthetic opioids are 2-benzyl benzimidazole compounds known as nitazenes. First identified in the US and Canadian drug supply in 2019, nitazenes are considered 10 to 40 times more potent than fentanyl (*5–8*). Furthermore, the molecular structures of synthetic opioids, including nitazenes, are consistently altered to skirt laws banning known compounds (*3*, *7*, *9*). While commercial tests exist for fentanyl, including lateral flow assay test strips (*10–12*), there is an immediate need for diagnostic assays to detect the range of nitazenes in the drug supply, as well as nitazene metabolic byproducts to assist medical providers with patient care (*13*).

Diagnostic assays often contain a biomolecule to specifically recognize the drug(s). To develop nitazene biosensors, we chose to redesign the binding pocket of the plant abscisic acid (ABA) receptor PYR1 (*14–16*). PYR1 binds a phosphatase HAB1 in the presence of ABA through an allosteric gate-latch lock mechanism (*17*), representing a natural chemical induced dimerization (CID) module. This CID module presents a built-in transduction mechanism coupling protein interactions to a diverse range of signal outputs (*16*, *18*, *19*). This is additionally advantageous because the molecular ratchet mechanism of the PYR1-HAB1 CID allows highly sensitive readouts from lower affinity ligand-receptor pairs (*14*, *20*). Immunoassays with antibodies or other binding molecules can be developed by brute force screening, but the relatively small surface area of drugs limits the affinity and ability to bind multiple related drug variants. While drug host receptors can be modified into a sensor (*21*), the same receptor can bind chemically dissimilar drugs, hampering unique identification of a drug class. Computational design has recently been used to create moderate affinity binders for largely apolar, rigid molecules (*22–26*). However, binders are not sensors: the transduction of the binding event into a measurable signal must typically be engineered for each bespoke design, limiting generality and throughput. In contrast, repurposing the PYR1 binding pocket, while keeping its transduction mechanism intact, promises to be a more generalizable solution.

In this study, we used computational design of the PYR1 binding pocket to identify biosensors suitable for detection of nitazenes and their byproducts. We used deep mutational scanning, directed evolution, and computational modeling to create a pan-nitazene sensor that can detect multiple nitazene derivatives, including the common variant isotonitazene and its 4-hydroxy nitazene byproduct. We then developed a luciferase-based *in vitro* diagnostic platform that is label-free, fast, sensitive, and can be performed in complex biological matrices like urine. The computational-experimental methodology represents a general way to rapidly develop biosensors to address the emergence of new synthetic opioids that can circumvent existing detection modalities.

## RESULTS

### A computational design protocol for sensing the nitazene family of synthetic opioids

Design of an allosteric biosensor requires solving the challenge of protein-ligand binder design consistent with, and constrained by, a structural definition of the transduction mechanism (*27*). A major advantage of redesigning PYR1 for new ligand sensing is that the receptor has an exceptionally well understood and characterized CID mechanism, where a bound water maintains hydrogen bonds between the ligand, the PYR1 receptor, and the HAB1 protein (*17*) (**Figure 1A,B**). We hypothesized that successful designs would recapitulate the spatial orientation of this ligand hydrogen bond acceptor.

**Fig. 1.**
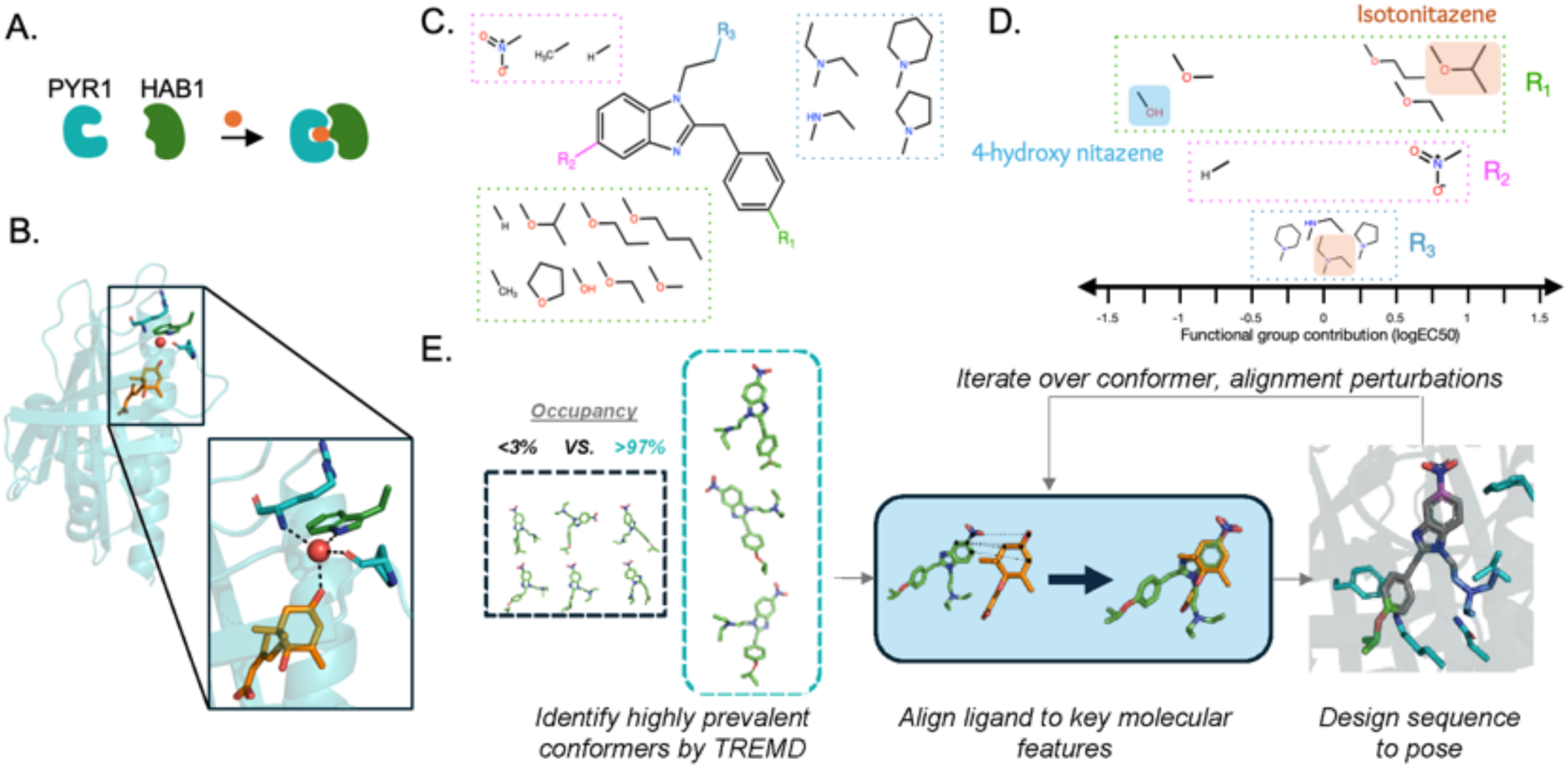
Computational design of a pan-nitazene biosensor. **(A)** Cartoon of the PYR-HAB chemically induced dimerization mechanism. **(B)** The structural definition of the transduction mechanism for PYR1 biosensors. The ligand shown as orange sticks is abscisic acid, the original PYR1 ligand. Red sphere shows the bound water molecule. (**C**) Structure of the nitazene central 2-benzyl benzimidazole with locations for possible substitutions color-coded. **(D)** Functional groups of nitazene derivatives organized by substitution position and contribution to molecular potency. The sign shows the direction of effect on potency (negative values are weaker potency), and the magnitude (logEC50) reflects contribution strength. Values are regression model coefficients from one-hot encoded functional group identities at each position. The R_1_ groups of isotonitazene and its less-potent metabolic product 4-hydroxy nitazene are labeled, as well as the common tertiary amine at R_3_ shared by both molecules. **(E)** Overview of the computational design process. Temperature replica exchange MD (TREMD) is used to identify highly prevalent solution populated ligand conformers. These conformers are aligned to key molecular features preserving the PYR1 transduction mechanism, and sequences are designed to each pose. This process is iterated over conformer and alignment perturbations and filtered to identify sets of sequences. A library is encoded using position-specific and local residue preferences.

To guide our design efforts, we first sought to understand the relationship between the molecular features of nitazenes and their potency. The nitazene family of 2-benzyl benzimidazole compounds contains substitutions observed at three positions (R_1_, R_2_, R_3_ ; **Figure 1C**) (*7*). We used supervised learning to classify the relative importance to potency of these substitutions (**Figure S1**) using a previous study of nitazene substitutions on cell-based activation of the μ-opioid receptor (*7*). A nitro group at R_2_ and larger aliphatic ethers at R_1_ increased potency significantly (**Figure 1D**), consistent with known potent synthetic opioids like isotonitazene (*7*, *28*), which contains an isopropoxy branched ether group at R_1_. Substitutions of different N-containing heterocycles or secondary and tertiary amine groups at R_3_ have less effects on molecular potency. Finally, isotonitazene and other nitazenes are metabolized at R_1_ to an hydroxyl (*7*, *29–31*), forming 4-hydroxy nitazene and related metabolites. This leaves the R_2_ group as a defining feature of nitazene potency. Thus, a biosensor that can detect a range of nitazenes containing an R_2_ nitro group and with limited cross-reactivity to other opioids is imperative for sensing nitazenes in the drug supply.

Repurposing an existing pocket to sense nitazenes requires a way to down-select the astronomical number of possible ligand-protein conformations. We accomplished this by developing a new computational design protocol to identify likely ligand-protein conformations and ligand orientations in the receptor binding pocket that maintain the transduction mechanism (**Figure 1B,E**). Sequences for these poses are then designed using either physically based (*32*) or deep learning (*33*) algorithms.

To identify likely conformers of isotonitazene, which contains 9 rotatable bonds, we performed temperature replica exchange MD simulations in solution. Three isotonitazene conformers were observed greater than 97% of the time (**Figure S2**). To orient these conformers in the binding pocket, we reasoned that likely configurations occur where the important nitro group hydrogen binds to the bound water critical to the dynamic transduction mechanism. Performing the alignment revealed that two isotonitazene conformers could dock without steric clashing when all 21 allowable positions in the binding pocket are mutated to glycine. These results match ligand docking experiments in a deeply mutagenized PYR1 library biased toward hydrophobic ligands (HMH, (*34*)), which contains sequences that allow docking of nitazenes in the binding pocket (**Figure S3)**.

To test these initial steps of our computational design process, we screened a nitazene panel (**Table S1**) against the HMH library using a yeast two hybrid (Y2H) assay (*35*, *36*), identifying sensors for six nitazene family members with differences in R_1_, R_2_, and R_3_ groups (although not isotonitazene; see **Figure S4, Table S1, Table S2**). Initial nitazene hits were identified with a geometric mean minimum dose response (MDR) of 61 µM in the Y2H growth selection, suggesting that their sequences were not optimized for binding. The hits had a mean of 6.4 mutations from the parental receptor; a previous computationally designed double site saturation mutagenesis library (*18*) yielded no sensors when nearly the same panel was screened (**Table S1**), highlighting that many PYR1 mutations are necessary to repurpose this privileged receptor for recognizing nitazenes.

### Computational design protocol yields a nM-responsive nitazene binder

The insights from screening and structural modeling were integrated into the design process for a nitazene-specific PYR1 library. We used the perturbed alignments of predicted conformations of isotonitazene for Rosetta sequence redesign (see **Methods, Table S3**), fixing the identity of residues known to be essential for the allosteric transduction mechanism (*17*, *35*). The library was ordered as an oligo pool and constructed by a four part Golden Gate assembly (*37*) in a thermally stabilized PYR1 background, PYR1^HOT5^ (*38*) (**Table S4-S5**, **Figure S5)**. The library contained a mean of 9 mutations from the PYR1^HOT5^ background. We screened this library using Y2H assays against a panel of nitazenes (**Figure 2A**), identifying 49 unique ligand-responsive sequences for six nitazene analogs, including isotonitazene (**Figure S6, Table S6**). Sensors with the lowest MDR for each compound are shown in **Figure 2B**. Across all hits, the MDR was 19 µM, representing a significant improvement over that of the previous HMH library (61 µM; two-sided Wilcoxon rank-sum test, p-value 8.5e-5).

**Fig. 2.**
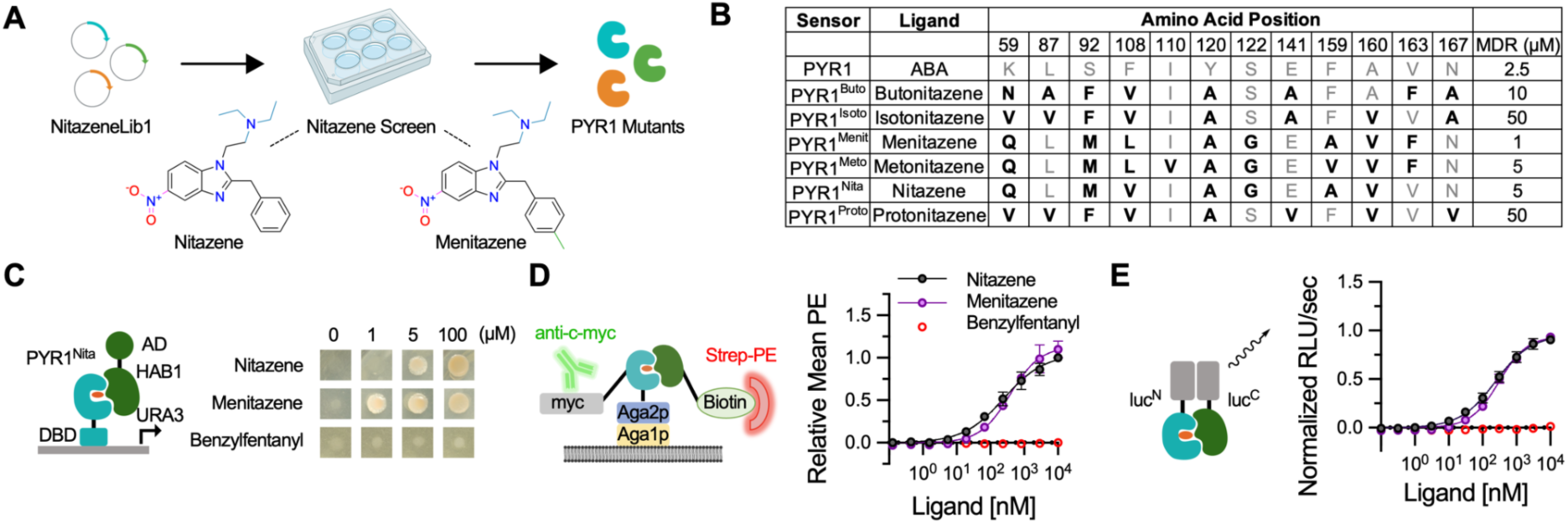
Isolation of PYR1^nita^, a nanomolar sensor for the synthetic opioid nitazene. **(A)** Schematic of the sensor isolation pipeline, including chemical structures of nitazene and menitazene. NitazeneLib1 was screened for sensors using Y2H growth selections in the presence of a ligand of interest. **(B)** Binding pocket mutations of PYR1 mutants hit from NitazeneLib1. The most sensitive receptor for each nitazene derivative is shown. Mutations from the wild type residues shown in bold. **(C-E)** PYR1^nita^ portability to different sensing modalities: Y2H growth assays (C), yeast surface display (D), and in vitro split nano-luciferase assays (E). (D,E) n=4, data points represent the mean, error bars represent 1 s.d. and in some cases may be smaller than the symbol.

From this screen, we identified PYR1^nita^ (PYR1^HOT5/H60P/N90S^ with K59Q, S92M, F108V, Y120A, S122G, F159A, A160V) as the lead candidate. This mutant appeared in screens for nitazene (MDR: 5 µM), menitazene (MDR: 1 µM), and protonitazene (MDR: 50 µM), and showed no detectable binding to the structurally unrelated synthetic opioid benzylfentanyl (**Figure 2C, Figure S6**). To further characterize PYR1^nita^, we displayed it on the surface of yeast and performed cell surface titrations. No binding was observed in the presence of a thermostable variant of HAB1 (ΔN-HAB1^T+^) alone or in the presence of the structurally unrelated ligand benzylfentanyl. Saturable responses were observed for nitazene EC_50_ 250 nM (95% c.i. 177-390 nM) and menitazene EC_50_ 383 nM (95% c.i. 277-588 nM), with a limit of detection (LOD) for nitazene of 43 nM (**Figure 2D**). To test binding *in vitro*, we optimized a previously described *ex vivo* split luciferase detection system (*18*) using a split Nanoluc (*39*). Here, LgBiT is fused to a PYR1 sensor construct, and SmBiT is fused to HAB1 (*40*) (**Figure S7**). Ligand titrations showed no activation with benzylfentanyl and nanomolar sensitivity to both nitazene EC_50_ 243 nM (95% c.i. 216-276 nM) and menitazene EC_50_ 306 nM (95% c.i. 240-405 nM), with a LOD for nitazene of 3.2 nM (**Figure 2E**). Thus, the computational design pipeline enabled the isolation of a nitazene sensor with low nM sensitivity *in vitro*.

### Optimization of nitazene biosensors by deep mutational scanning

Next, we used deep mutational scanning coupled with a directed evolution strategy to develop two distinct nitazene biosensors (**Figure 3A**). The first of these is for the detection of 4-hydroxy nitazene, the major metabolite byproduct of isotonitazene. While the current needed LOD is unknown (*41*), an effective sensor should exhibit a low nM LOD and function in complex biological matrices like urine. The second sensor is a pan-sensor capable of detecting nitazenes in mixtures. While the sensitivity requirement of this sensor is less strenuous, it must be specific for nitazenes compared with other synthetic opioids, fillers, or recreational drugs.

**Fig. 3.**
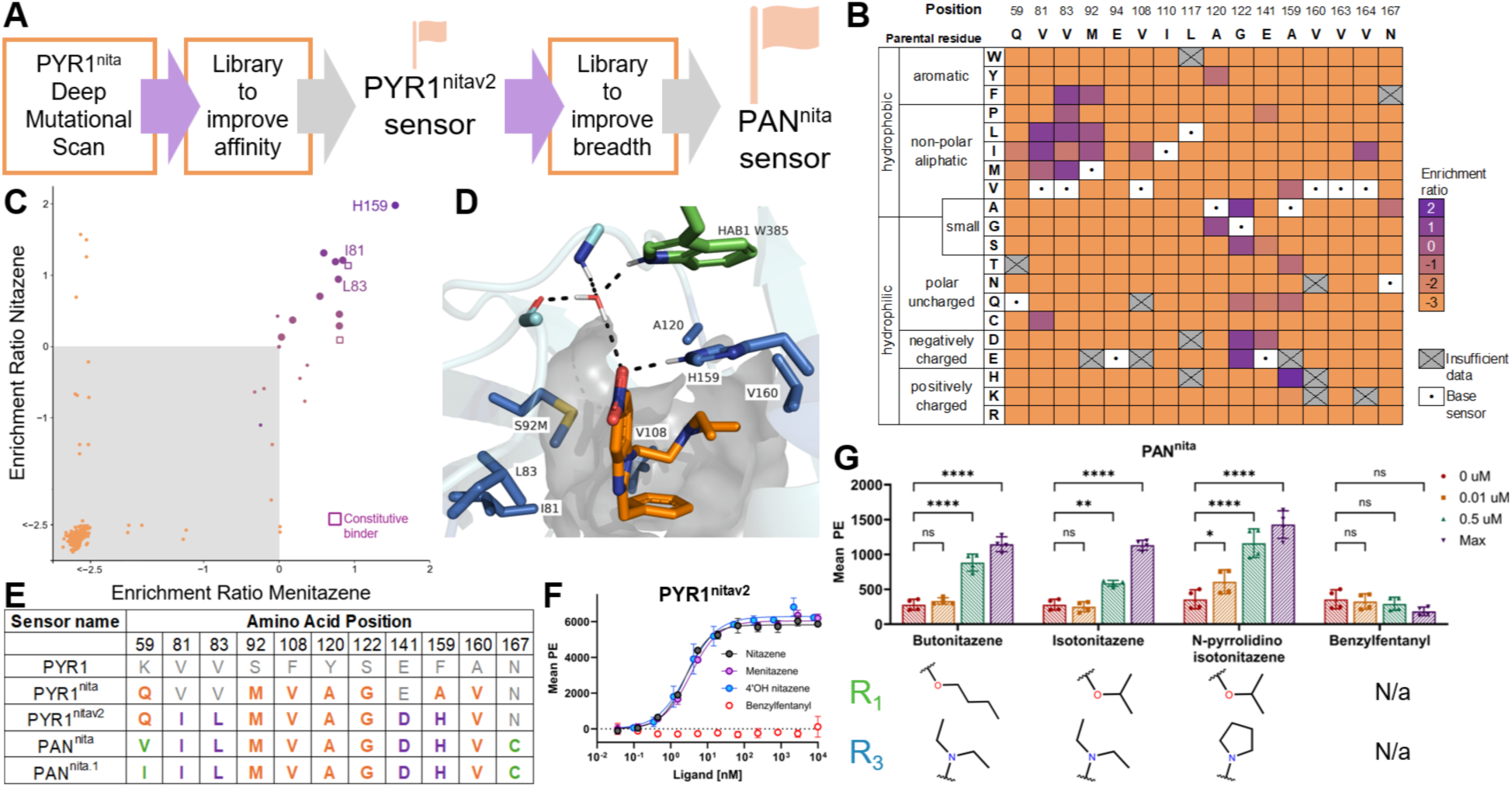
Optimization of nitazene sensor breadth and sensitivity using deep mutational scanning. **(A)** Overview of the protein engineering workflow. **(B)** Heatmap of calculated enrichment ratios of single-point mutations to the PYR1^nita^ sensor in binding nitazene. 16 residues relevant to ligand binding were scanned. Purple indicates an increased enrichment ratio and orange indicates a decreased enrichment ratio relative to the parental sensor sequence. **(C)** Correlation plot of the enrichment ratios of PYR1^nita^ binding nitazene versus menitazene. The color scale correlates to that of the heatmap. Open purple squares indicate mutations that result in constitutive binding. **(D)** Model of the PYR1^nitav2^ computationally designed structure. Structure of the PYR1^nitav2^ binding pocket, highlighting the histidine residue at position 159 which is hypothesized to delocalize the partial charge on the nitro group. Selected mutated residues from PYR1 are shown as cornflower blue sticks. **(E)** Sequence differences between WT PYR1, computationally designed PYR1^nita^, PYR1^nitav2^, and PAN^nita^. New mutations added in each step of the optimization are shown in orange, purple, and green, respectively. **(F)** Yeast surface display titrations of PYR1^nitav2^ against nitazene (black circles), menitazene (purple circles), 4-hydroxy nitazene (cornflower blue circles), and benzylfentanyl (open red circles). The EC_50_ was 2.5 nM (95% c.i. 2.2 to 2.9 nM), 3.4 nM (95% c.i. 3.0 to 4.0 nM), and 2.9 nM (95% c.i. 2.2 to 3.9 nM) for nitazene, menitazene, and 4-hydroxy nitazene, respectively, using 500 nM of biotinylated ΔN-HAB1^T+^. **(G)** Yeast surface display binding measured for PAN^nita^ against indicated ligands. Max concentrations for butonitazene, isotonitazene, and N-pyrrolidino isotonitazene are 24 uM and 10 uM for benzylfentanyl. R1 and R3 groups that differ between nitazenes are indicated. **** p<0.0001, ** p<0.002, * p<0.05, ns not statistically significant. For all measurements, n=4 comprising two technical replicates performed on different days. Error bars represent 1 s.d.

In a first step to designing these sensors, we performed a deep mutational scan (DMS) of 16 positions inside of the PYR1^nita^ binding pocket using yeast display screening coupled to fluorescence activated cell sorting. Libraries were screened at 250 nM of nitazene and menitazene (approx. 20% of the EC_50_). Additionally, a constitutive (no ligand but in the presence of the 500 nM HAB1^T+^ binding protein) and a reference control (sorting on all gates except the gate associated with binding) were included. Example sorting gates are shown in **Figure S8**.

Following deep sequencing of these populations, the frequencies of each mutant were assessed and normalized to the parental sequence by an enrichment ratio (**Figure 3B, Figure S9**). Most mutations (298/320) were depleted, indicating a restricted sensor pocket. Nitazene and menitazene shared 10 non-constitutive beneficial mutations (V81CLI, V83ML, M92FL, A120G, A159H, V164I; **Figure 3C**), largely in the aliphatic central cavity near the predicted binding site of the benzylimidazole nitazene scaffold (**Figure 3D**). The largest enriched mutation for both ligands was A159H. Our original design models did not include a residue which delocalizes the partial charge on the nitro group; we hypothesize that H159 satisfies this requirement (**Figure 3D**). Indeed, redesigning the PYR1 sequence using the deep learning algorithm LigandMPNN (*33*) identifies H159 in 8% of designs **(Figure S10)**. Overall, the mutational profile is largely consistent with the designed binding mode.

To develop a sensor capable of binding the major metabolite byproduct 4-hydroxy nitazene, we used the DMS output to create a focused combinatorial library of 4,608 members largely containing the beneficial mutations shared between nitazene and menitazene (**Table S7**). Screening this library on nitazene and menitazene yielded two sensors, PYR1^nitav2.1^ (PYR1^nita^ with V81I, M92F, A120G, G122E, E141D, A159H) and PYR1^nitav2^ (PYR1^nita^ with V81I, V83L, E141D, A159H), containing 4-6 additional mutations from PYR1^nita^ (**Figure 3E** and **S10**). Both sensors recognized 4-hydroxy nitazene, menitazene, and nitazene at an average LOD of 100 pM and EC_50_ of approx. 2 nM (**Figure 3F**, and **S11)**, representing a more than 100-fold improvement in affinity over the originally designed PYR1^nita^ sensor. Neither sensor recognized the unrelated synthetic opioid benzylfentanyl at the highest concentration tested, indicating highly specific recognition of nitazene, its metabolic byproduct 4-hydroxy nitazene and a close analog menitazene over a competing synthetic opioid.

To develop a sequence able to sense structurally diverse nitazenes, we used the structural model of PYR1^nita^ to identify mutations which could allow binding breadth by increasing the pocket space for diverse functional R2 and R3 groups. We created a focused combinatorial library of 2048 variants encoding differences at positions 59, 141, 163, 164, & 167 to the PYR1^nitav2.1^ and PYR1^nitav2^ sensors (**Table S8**). After the library was screened against constitutive binders, the library was split and screened in parallel against butonitazene, isotonitazene, and N-pyrrolidino isotonitazene (**Figure S12**). These populations were deep sequenced; sequences that were enriched in all three ligand populations were individually tested (**Figure S13)**. The best sensors, PAN^nita^ and PAN^nita.1^, recognized all three diverse nitazenes and did not recognize benzylfentanyl (**Figure 3G** and **S14**). PAN^nita^ has in total 11 mutations in the receptor binding pocket out of 18 total mutable positions. Combined, computational design coupled to protein engineering can access a more diverse functional sequence space than accessed by previous libraries.

### Development of a robust diagnostic assay suitable for biological matrices

To test whether PYR^nitav2^ and PAN^nita^ are capable of sensing nitazenes under relevant conditions, we repurposed our *in vitro* luminescence assay. Luminescence detection assays are susceptible to errors from varying experimental conditions including fluctuating activity in diverse biological matrices (*42*). For these reasons, ease of single sample detection is limited by the need for external calibration. A previously described calibrator luciferase (*43*) enables single-sample ratiometric quantitative readouts **(Figure 4A)**. This assay compares the ratio of ligand dependent luminescence to background GFP fluorescence to mitigate experimental variance. We tested this new calibrator assay with the parental PYR1 sensor and a range of synthetic cannabinoid sensors (*18*). All tested sensors had nearly identical dynamic range, LOD, and EC_50_ values between buffer and urine **(Figure S15)**. Using this assay, PYR1^nitav2^ could recognize 4-hydroxy nitazene with a LOD of 1 nM in urine, and with no cross-reactivity to the unrelated opioids benzylfentanyl, codeine, or heroin (**Figure 4B**). We next tested whether PAN^nita.1^ could recognize diverse nitazenes relative to these other synthetic opioids. PAN^nita.1^ bound the nitazene variants butonitazene, isotonitazene, and N-pyrrolidino nitazene while exhibiting minimal perception of benzylfentanyl, codeine, and heroin. At their highest tested concentration, this assay was able to detect a 6.4x, 7.2x, and 23x fold-change respectively for the target ligands butonitazene, isotonitazene, and N-pyrrolidino nitazene, with limited activation of off-target opioids (2-way ANOVA, p<0.0001) (**Figure 4C**). Similar results were obtained for the PAN^nita^ sensor (**Figure S16**).

**Fig. 4.**
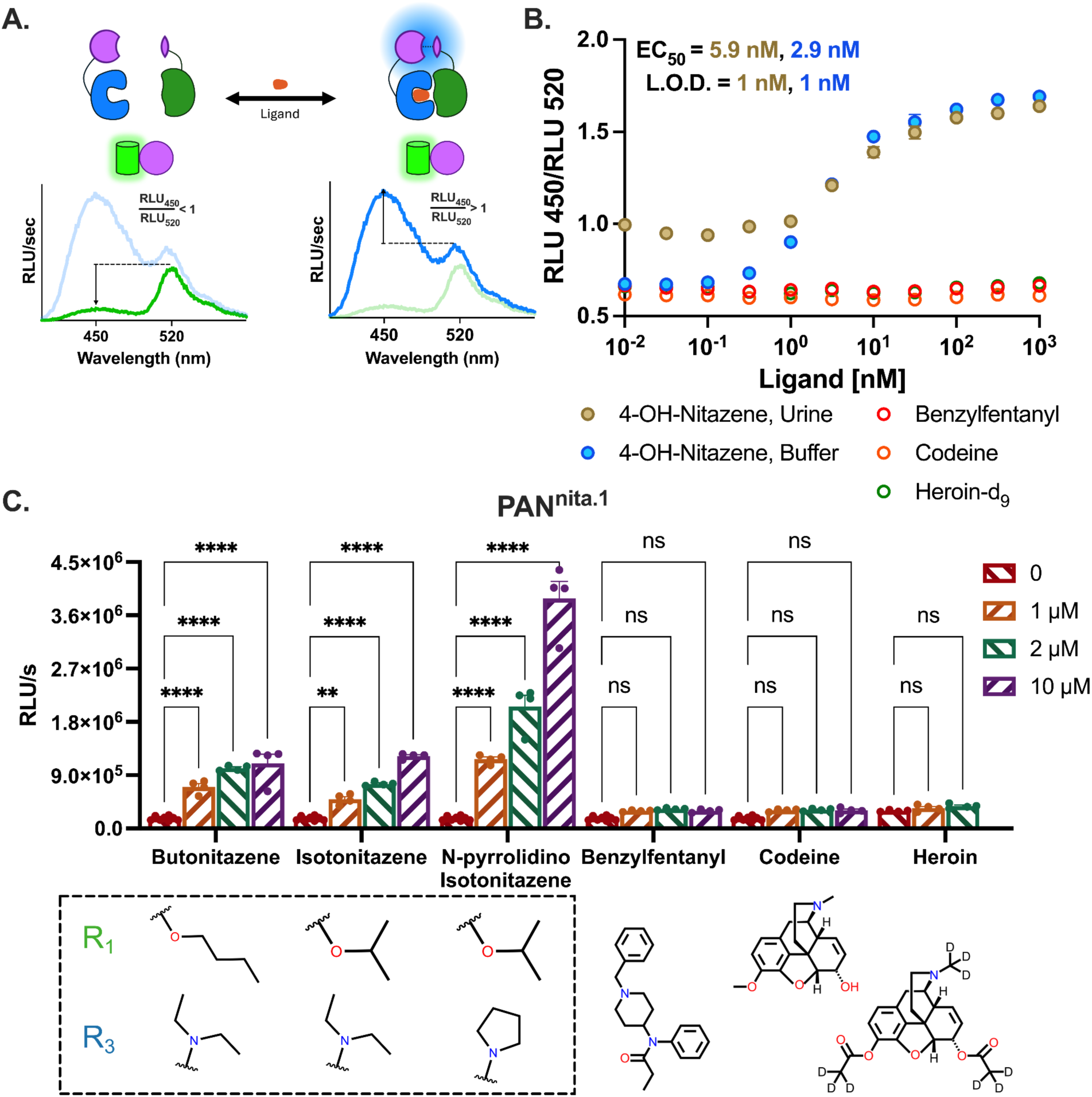
A ratiometric luminescent assay can sense nitazenes specifically in clinically relevant matrices. **(A)** Cartoon of the luminescence assay. A mNeonGreen-NanoBiT “calibrator” protein enables ratiometric detection of samples, fluorescing at a wavelength of 520 nm independent of ligand concentration. PYR1 to ΔN-HAB1^T+^ dimerization results in an increased ratio of the relative luminescence units per second (RLU/s) at 450 nm versus the RLU/s at 520 nm. **(B)** Ratiometric assay of PYR1^nitav2^ in buffer or in urine using the indicated ligands. EC50 measurements and LOD for the sensor are colored for the appropriate condition. 200 nM SmBit-ΔNHAB1, 10 nM LgBiT-PYR1, and 512 pM calibrator are used. Benzylfentanyl, codeine, and heroin were tested in buffer. **(C)** Luminescence assay results using PAN^nita.1^ show sensitivity against a panel of nitazene variants and synthetic opioids across multiple concentrations. Analysis of variance was calculated using an ordinary 2-way ANOVA and Dunnett’s multiple comparison test, with a single pooled variance (**** p<0.0001, ** p<0.01, ns not statistically significant). 4 nM SmBit-ΔNHAB1 and 4 nM LgBiT-PYR1 are used. Data is shown as the average of n=3 (A,B) or n=4 (C) replicates. Error bars represent the standard error of the mean and in some cases are smaller than the symbols. LODs were calculated using the 3σ method, which is equivalent to the drug concentration that yields a signal equal to 3 times the standard deviation of the blank after subtraction.

## DISCUSSION

Computational protein design has advanced towards relevant and pressing societal needs (*44*). Here we designed and engineered protein biosensors for a range of clinically relevant synthetic opioids. The designed sensors, the best of which exhibit pM responsiveness and low nM EC_50_s, are orders of magnitude more sensitive than has been achieved by previously described computational approaches for small molecule binders (*22*, *24*, *25*). Our design success depended on a quantitative, molecular understanding of the dynamic transduction mechanism, which was worked out by biochemical and structural studies of abscisic acid perception in plants (*17*, *35*, *45–48*). A deeper mechanistic insight into the more complicated and varied allosteric transduction mechanisms of ligand-dependent molecules like GPCRs or bacterial transcription regulators (*27*) could improve biosensor hit rates for these classes of proteins.

Variants of the PYR1 receptor are able to sense an unusually broad spectrum of drug-like molecules (Tian et al, unpublished). This suggests that PYR1 sensors can be developed for a wide range of molecules, ultimately enabling new medical diagnostics, chemically-responsive cell therapies, environmental sensors (*19*), and biotechnologies for cell engineering (*34*). Our design process allows navigation to a search space largely inaccessible to random site saturation mutagenesis libraries, and when coupled with our conformer selection and alignment protocol creates a powerful design approach for ligand binding. We anticipate that emerging (*49*, *50*) and future deep learning algorithms will further improve design and overall hit rates in designing PYR1 receptors for new target ligands, which could be incorporated into our protocol in the future as conformer selection and alignment are largely independent of sequence generation. Another possible approach to develop biosensors for wider swathes of chemical space is the generation of de novo proteins which maintain the PYR1 dynamic transduction mechanism. These efforts would be enabled by new computational and experimental tools to test and predict motions of designed proteins (*51*, *52*).

## Acknowledgments

We would like to thank Yves Janin for kindly providing the luciferase prosubstrate, Hikarazine-108. ACL would like to thank the Interdisciplinary Quantitative Biology and Molecular Biophysics programs at the University of Colorado Boulder for ongoing support. TAW would like to thank M. Stammnitz for helpful discussions related to transduction mechanisms for ligand-dependent protein biosensors.

## Funding

National Science Foundation NSF Award #2128287 (TAW)

National Science Foundation NSF Award #2128016 (SRC, IW)

National Science Foundation NRT Integrated Data Science Fellowship Award #2022138 (LMW)

NSF GRFP Award #1650115 (ACL)

NSF GRFP Award #2040434 (ZTB)

DARPA CERES Award#D24AC00011-05 (SRC, IW, TAW)

NIH award# R01GM123296 (LMW, AJF)

NIH award #5T32GM145437 (LMW)

## Author contributions

Non co-1st author trainees are listed in alphabetical order.

Conceptualization: ACL, CLM, NRR, IW, TAW

Methodology: ACL, CLM, RC, SS, RD, AJF, MRS, IW, TAW

Investigation: ACL, CLM, RC, SS, ZTB, RD, AJF, NRR, NS, JW, LMW

Visualization: ACL, CLM, RC, SS, IW, TAW

Funding acquisition: ACL, ZTB, SC, MRS, IW, TAW

Project administration: IW, TAW

Supervision: SC, MRS, IW, TAW

Writing – original draft: ACL, CLM, RC, SS, IW, TAW

Writing – review & editing: SC, MRS, IW, TAW

## Competing interests

TAW, SRC, and IW, have filed a provisional patent entitled REAGENTS AND SYSTEMS FOR GENERATING BIOSENSORS (US9738902B2; WO2011139798A2) covering some research in the present work. TAW is a consultant for Inari Ag and serves on the scientific advisory board for Metaphore Biotechnologies and Alta Tech.

## Data and materials availability

An automated protocol for the TREMD conformer generation protocol is available on GitHub https://github.com/ajfriedman22/SM_ConfGen. PyRosetta scripts for biosensor design by structural replacement are available at https://github.com/alisoncleonard/Structural-Replacement-Biosensor-Design. Scripts used to generate figures S1 and S10 are available at https://github.com/WhiteheadGroup/Leonard_ComputationalDesign_Supplemental.Raw deep sequencing data are deposited in the SRA (BioProject ID PRJNA1256820), and analyzed deep sequencing data are on Zenodo (doi:10.5281/zenodo.15298585). All other data are available in the main text or the supplementary materials.

## Supplementary Materials

### Materials and Methods

#### Chemicals

Nitazenes and other pharmaceutically active molecules for use in yeast surface display and split luciferase assays were purchased through Cayman Chemical, including 4’ hydroxy nitazene (#30218), benzyl fentanyl (hydrochloride, #19883), butonitazene (exempt preparation, #38653), codeine (CRM, #ISO60140), ethyleneoxynitazene (citrate, #34947), heroin-d9 (exempt preparation, #42070), isotonitazene (CRM, #30880), menitazene (citrate, #36634), metonitazene (citrate, CRM, #40808), N-desethyl etonitazene (#32512), nitazene (citrate, #36635), N-pyrrolidino isotonitazene (citrate, #34909), and protonitazene (hydrochloride, CRM, #38143).

#### Computational Design Protocol

The mutational library NitazeneLib1 was generated using a structural replacement computational docking protocol to generate high-quality plausible protein-ligand alignments of simulated low-energy ligand conformers.

To generate ligand conformers by molecular dynamics, an initial 3D conformer was downloaded from Pubchem (*1*). Conformers were generated by temperature replica exchange molecular dymanics simulations (TREMD) exactly as described in Leonard et al (*2*). For this paper, we developed an automated protocol with open source code (https://github.com/ajfriedman22/SM_ConfGen). We used the default implemented settings which run 27 temperature replicas ranging from 300 to 450 K and all simulations were run for 100 ns with replica exchange occurring every 2 ps. The conformer with the highest observed occupancy in the lowest temperature replica was given a relative free energy of 0.0 kcal/mol, and the free energy of other conformers were calculated relative to the highest observed occupancy assuming a Boltzmann weighted ensemble. Generated conformers were plotted sequentially by free energy relative to the lowest energy conformer, categorizing conformers as either “high occupancy” (greater than 4% occupancy cutoff used for menitazene and isotonitazene) or “low occupancy” (less than 1% occupancy cutoff used for menitazene and isotonitazene) (**Figure S2**). Low-energy conformers only were exported as a single sdf file for ligand docking.

Low-energy ligand conformers generated by TREMD were docked into the PYR1 protein ligand binding pocket using the “dock to sequence” approach of our method “Structural replacement for the design of protein - small molecule binding”, available as custom python scripts at https://github.com/alisoncleonard/Structural-Replacement-Biosensor-Design. In the dock to sequence method, the wild-type protein sequence to design is mutated to match the sequence of a known low-affinity ligand binding sequence identified through library screening in yeast two-hybrid (**Figure S4**), enabling visualization of ligand alignments that could accommodate the known binding sequence. Each nitazene conformer was docked in space using a Singular Value Decomposition (SVD)-based superposition method to align the 3D dimensional coordinates of a given novel query ligand with a target molecule. To engineer PYR1, the PDB structure 3QN1 (*3*) was used as the template, with atoms surrounding the conserved nitro group at nitazene position R_2_ superimposed on the target molecule abscisic acid carbonyl, with the nitro group acting as the required hydrogen bond acceptor needed for binding (*4*).

The alignment process begins with two sets of atomic coordinates: one from the query molecule to be aligned and the other from the target molecule. These do not need to be the full set of atomic coordinates for the molecules but must include at least three atoms that are not collinear. Each coordinate set is arranged into a matrix, where each row in the matrix is the x, y, and z coordinates of an atom, and the corresponding row in the other matrix is the corresponding atom in the other molecule. To obtain a set of transformation matrices, each coordinate set is translated so that its center of mass (mean) is positioned at the origin, then a covariance matrix is calculated between the two centered coordinate sets, and finally the covariance matrix is decomposed into three components: two orthogonal matrices and a diagonal matrix of singular values which are used to calculate a rotation matrix. Once the rotation matrix is computed, the original query compound undergoes transformations applied using the apply_transform_Rx_plus_v function available in PyRosetta, where the query coordinates are first translated so that their center of mass is at the origin, to ensure that the rotation is applied correctly, next the computed rotation matrix is applied to the coordinates of the query compound, to put it in plane with the target molecule, and finally the query is translated to the center of mass of the target molecule coordinates, completing the alignment.

As it is unlikely that a novel ligand would dock in exactly the same orientation as abscisic acid, the aligned ligand was next perturbed in space using the PyRosetta Rigid Body Perturbation mover. The moved ligand was verified both to not clash with the protein backbone coordinates, determined using a fast-grid search of PYR1 subjected to a poly-glycine shave, and to contain a nitrogen or oxygen close enough to the latch water molecule of the PYR1 structure to form a hydrogen bond. If these conditions were satisfied, the PYR1-ligand complex was relaxed and repacked using the PyRosetta Packer function, and the repacked complex was checked for collisions with the full protein including both backbone and side chains by fast-grd search. Finally, complexes that pass clash check were scored using the standard PyRosetta scorefunction, with complexes below −300 REU generally considered to be realistic protein-ligand complexes. This process can be repeated as many times as desired to appropriately map plausible ligand alignments in the protein pocket (**Table S2**).

Modeled protein-ligand structures were used as inputs for Rosetta FastDesign (*5*) (scripts available at https://github.com/alisoncleonard/Structural-Replacement-Biosensor-Design) for structure-informed rational design to select mutations for inclusion in the library NitazeneLib1. The results presented in Figure S10 were a LigandMPNN (*6*) sequence redesign of a similar set of modeled protein-ligand structures using (https://github.com/dauparas/LigandMPNN).

#### Plasmids and plasmid library generation

A full list of primer sets (**Table S9**) and plasmids (**Table S10**) used in the work are included. All plasmids were sequence verified using Oxford NanoPore (Plasmidsaurus).

To construct NitazeneLib1, libraries containing desired combinations of mutations were encoded in oligo pools (Agilent) divided into 3 cassettes (**Table S4**), along with 2 constant region cassettes flanking the PYR1 gene, to be cloned into the PYR1 yeast two-hybrid destination vector pND005 (*7*). Three sub-libraries were prepared, with each containing 2 of the 3 variable portions and one constant cassette (e.g. Library A contained variable cassettes 1 and 2, plus a constant cassette 3). Golden Gate assembly, transformation into *E. coli* XL1-Blue, and preparation of plasmid DNA were exactly as previously described (*7*). Transformation efficiencies (**Table S5**) were estimated using serial dilution, and library quality (**Figure S5**) determined by short read deep sequencing (Azenta Amplicon-EZ). PYR1^nita^ was assembled into pND003 to make pRMC075 by using Golden Gate assembly and it was transformed into EBY100 *S. cerevisiae* cells as previously described (*8*).

A single site saturation mutagenesis (SSM) library at 16 positions (59, 81, 83, 92, 94, 108, 110, 117, 120, 122, 141, 159, 160, 163, 164, and 167) was prepared for PYR1^nita^ (PYR1^HOT5/H60P/N90S/K59Q/S92M/F108V/Y120A/S122G/F159A/A160V^) by nicking mutagenesis (*9*) using pRMC075 as a plasmid template. NNK mutagenic primers were designed using published SUNi mutagenesis scripts (*10*). Cells were transformed into XL1-Blue electrocompetent *E. coli*. Plasmid DNA was extracted with a ZymoPURE II Plasmid midiprep kit. Transformation of the library into EBY100 *S. cerevisiae* cells was performed as previously described (*8*).

A pan-nitazene library (Nitazene Library 2, **Table S7**) of 4608 members was made with mutations encoded into ultramers (IDT) with constrained degenerate mixed base codons and assembled according to (*7*). A single forward ultramer (Nitz_Combo_Lib2_for) spanning residues 58 to 119 and encoding V81VIL (VTT), V83VILM (VTK), M92MILF (WTK), V108VI (RTH) was amplified with a mix of two reverse ultramers (Nitz_Combo_Lib2_rev_159AV, Nitz_Combo_Lib2_rev_159H) spanning residues 109-168 and encoding A120GA (GSS), G122DEAG (GVN), E141DE (GAW), and either A159GA (GYG) or A159H (CAT) generating a full length Golden Gate ready cassette. A single gBlock (IDT) (Nitz_ComboLib2_Block_1_3) encoding both N-terminal sequence (before residue 58) and C-terminal sequence (after residue 168) with appropriate Golden Gate cut sites was used to generate the full length CDS in the library with pND003 destination vector. The library was prepared for YSD as described above. PYR1^nitav2^ and PYR1^nitav2.1^ were isolated as colonies to obtain plasmids P5 and P8 after sorting.

Two libraries based off of PYR1^nitav2^ and PYR1^nitav2.1^ were constructed via combinatorial mutagenesis essentially according to (*15*) to make Nitazene Library 3 and Nitazene Library 4 (**Table S8**), respectively. Each library had the following mutations added by long ultramers (IDT) (pZB717-pZB720) encoding Q59QVASLIMT (DYR), D141EDAG (GVN), A159VA (GYT) or A159H, V163VA (GYN), V164VILM (VTK), and N167SACGN (KSY), where A159VA and A159H were encoded on two separate ultramers, for a total of 2560 library members each. After Nitazene Libraries 3 and 4 were prepared for YSD as described above, they were pooled together to create Nitazene Library 5 containing 5120 library members total, which was screened according to conditions listed below. NGS was performed as described below to identify pan-nitazene contenders. Select individuals were cloned into pND003 with constant cassettes cass1_pND003_ACL and cass5_pND003_ACL and gBlocks (IDT) containing the corresponding mutations for each contender 1_Cas_2-4_Nit_pan_contender through 6_Cas_2-4_Nit_pan_contender were created via Golden Gate assembly according to Daffern et al., 2023 (*11*).

pYTK047 was a gift from John Dueber (Addgene plasmid # 65154; http://n2t.net/addgene:65154; RRID:Addgene_65154)

#### Yeast Two-Hybrid Screening of Mutant Biosensor Libraries

Selection experiments for mutant receptors that respond to new ligands were conducted as previously described by Beltrán (*4*). Briefly, the indicated pBD-PYR1 mutant library was transformed into MAV99 (MATa trp1-901 leu2-3,112 hisΔ200 ade2-101 gal4Δ gal80Δ can1rcyh2rLYS2::(GAL1::HIS3) GAL1::lacZ SPO13::10xGAL4site::URA3) harboring pAD-HAB1. A minimum transformation efficiency of 1 million CFU was achieved, typically by pooling two transformations. After transformations, negative selections were conducted twice to remove receptors that bind HAB1 in a ligand-independent fashion (i.e., constitutive receptors) by growing the library on solid media containing 1 g/L 5-FOA; the purged library was collected and stored as 30 (v/v) % glycerol stocks until use (*4*).

These glycerol stocks were used in subsequent selections for cells responsive to 100 μM nitazene ligands described in **Table S1** on SD-Trp,-Leu,-Ura media. Colonies supporting uracil-independent growth at 30 °C were isolated after 3 days, regrown on SD-Trp,-Leu,+1 g/L 5-FOA to screen out constitutive biosensors, retested to confirm ligand-dependent growth on SD-Trp,-Leu,-Ura plates with and without test chemical. The minimum dose response was determined by growth validation after 3 days, in which yeast cells were grown as spot assays by suspending a colony in 100 µL of 1X TE and dropping 2 µL onto SD-Trp,-Leu,-Ura plates containing various concentrations of the target ligand.

#### Library sorting by yeast surface display

Sorting of the PYR1 yeast displayed libraries was performed as done in Steiner et al.(*12*) except for the following differences. Cells were induced in SD/GCAA media containing a 9:1 ratio of galactose:dextrose. Induction was carried out at 22°C and 250 rpm for 20 hours. Cells were washed in citrate-buffered saline (20 mM trisodium citrate dihydrate, 147 mM NaCl, 4.5 mM KCl) supplemented with 0.1% w/v BSA adjusted to pH 8.0 with 1 M NaOH (CBSF), while prepared ΔN-HAB1^T+^ from plasmid pJS723(*4*) was resuspended in CBSF plus 1 mM DTT and 5 mM TCEP.

For the SSM library sort, each labeling condition had 250 µL of OD_600_ = 2 cells, 3 µL of ligand dissolved in DMSO for an overall solvent of 1 v/v%, ΔN-HAB1^T+^ to 500 nM, and the volume of the reaction brought to 300 µL with CBSF. Binding and labeling were performed as in Leonard et al.(*2*). Cells were collected at approximately half of EC_50_ in a diagonal gate, excluding background, selecting 200,000 cells each of the top 10-15% of binders, a reference population, and non-binders. Cells were grown and stocks prepared according to (*2*). Next generation sequencing (NGS) was performed on these screened cells as described below.

Nitazene Library 2 was screened with 250 µl of OD_600_ = 2 cells and maintained solvent at 1 v/v%, 100 nM of ΔN-HAB1^T+^, and a total volume of 300 µl attained by adding CBSF in order to obtain PYR1^nitav2^ sensors. First, 200,000 cells of the top 10% of binders were sorted using 250 nM menitazene. The second sort used identical reaction conditions, but only 200,000 cells of the top 5% of binders were collected. The final sort for the PYR1^nitav2^ sensors differed from the previous sort only by using 100 nM menitazene, which yielded PYR1^nitav2^ and PYR1^nitav2.1^ sensors, isolated as individual colonies. DNA was extracted from PYR1^nitav2^ and PYR1^nitav2.1^ and plasmid sequenced (Plasmidsaurus) to identify sensor sequences.

A total of 250 µl of OD_600_ = 2 cells of the combinatorial mutagenesis libraries of PYR1^nitav2^ and PYR1^nitav2.1^ was screened with 100 nM ΔN-HAB1^T+^ and the first sort was performed with no ligand and 1% (v/v) DMSO to collect 200,000 cells that did not bind constitutively. The resulting cells were grown and screened against three ligands in parallel, butonitazene, isotonitazene, and N-pyrrolidino isotonitazene at 2.5 µM, 2.5 µM, and 23.8 µM, respectively, where butonitazene and isotonitazene were dissolved in methanol and samples run maintained at 1% v/v% of solvent. The top 5% of binders were collected to get 200,000 cells and DNA was prepared for NGS as described below.

The top performing contenders for the PAN^nita^ sensor were identified as those that had the highest enrichment ratios for each ligand and gBlocks cassettes containing the identified mutations were cloned into pND003 via Golden Gate assembly to make pRMC079-084. These sensors were tested with yeast surface display with 100 nM ΔN-HAB1^T+^ and various concentrations of butonitazene, isotonitazene, N-pyrrolidino isotonitazene, and benzylfentanyl, maintaining a solvent concentration of 1% v/v.

#### Next generation sequencing of collected cell populations

DNA was extracted as described in Medina-Cucurella & Whitehead (*8*), except for the following differences. Instead of utilizing a Qiagen mini-prep column, an NEB miniprep column was utilized and DNA was washed with 550 µL DNA wash buffer two times and eluted in 30 µL nuclease free water. Product from the purification of plasmid with exonuclease I and lambda nuclease was cleaned using an NEB DNA & PCR cleanup kit according to the manufacturer’s instructions.

Plasmid DNA was prepared for NGS by amplifying with either primer sets RMC-P1035 and RMC-P1036 (length of 383 bp) or RMC-P1037 and RMC-P1038 (length of 388 bp). 10 µL of plasmid DNA was amplified using 12.5 µL Q5® High-Fidelity 2X Master Mix, 1.25 µL 10 µM forward primer, 1.25 µL 10 µM reverse primer, and 25 µL nuclease free water (NFW) and placed in the thermocycler with the following cycles: 30 s at 98°C, 18 cycles of 5 s at 98°C, 20 s at 57.5°C, and 30 s at 72°C, followed by a final extension of 2 minutes at 72°C and a hold at 10°C. PCR product was purified using an NEB Monarch DNA Gel Extraction kit. DNA was pooled for nitazene and menitazene top binders, then reference populations, and non-binding populations at equal masses and sent to Quintara for Amplicon Express next generation sequencing. Resulting sequences were merged with FLASh (*13*), filtered for appropriate read length and amino acid start sequence, and heat maps of calculated enrichment ratios were created as in Medina-Cucurella & Whitehead, 2018 (*8*).

#### Protein expression and purification

Proteins were expressed in *E. coli* BL21(DE3) and purified by Ni-NTA affinity chromatography exactly as described (*4*).

#### Split Luciferase *in vitro* Assay

Proteins, stored as ammonium sulfate precipitates, were pelleted at 17,000xg for 10 minutes and then resuspended in desalting buffer (50 mM HEPES-KOH, 10% w/v glycerol, 200 mM KCl, 1 mM DTT, 5 mM TCEP, pH 8.0). A 7K MWCO Zeba™ spin desalting column (ThermoFisher, Cat. No. 89882) was equilibrated with desalting buffer and then used to buffer equilibrate the resuspended protein, which was then stored on ice until use. Ligand was diluted to 10x working concentration in desalting buffer and 20 𝜇L of this dilution was added to the bottom of Greiner™ 96-well, white, flat bottom plate (Cat. No. 655075). In a separate tube, desired proteins are combined with the luciferase substrate Hikarazine-108 (a gift from Dr. Yves Janin, Centre National de la Recherche Scientifique) and diluted with desalting buffer with reducing agents to reach a final volume of 180 uL per well. For bioassay measurements, urine was added to this mixture such that it reaches a final concentration of 20% v/v. Note, add the substrate, Hikarazine-108, last, just before adding the mixture to the wells to minimize its depletion before the reaction has begun in the plate. This 180 μL mixture was scaled to encompass all wells and replicates. Using a multichannel pipette, 180 μL of this mixture are added to each well. The wells are mixed up and down using the pipette, not aspirating to avoid bubbles. The plate was then covered and placed on a plate shaker at 25°C and 750 rpm. After 30 minutes, the foil was removed, and a light mist of ethanol can be used to pop any bubbles that might have been introduced. The plate was placed on a plate reader and an autogain feature was used to adjust the gain at a wavelength of 450 nm for the brightest anticipated well, so that overflow does not occur. Here, we use a BioTek Synergy™ H1 Microplate Reader with an integration time of 1 second and a read height of 7 mm. The plate was then run, and data was output in units of relative luminescence units (RLU) per second.

For ratiometric detection, inclusion of the calibrator luciferase includes mNeonGreen fused to the full length NanoBiT luciferase, enabling bioluminescence resonance energy transfer (BRET) between the pair (*16*). This protein sequence was ordered as an eBlock from IDT **(Table S11)** and cloned into a protein expression backbone from pJS723 using Gibson Assembly (*17,18*). The NanoBiT converts luciferase prosubstrate into blue light (λ_max_∼450 nm) while mNeonGreen gives off green light (λ_max_∼520 nm) and the optimized system yields two distinct spectrum peaks. Complex formation is measured as the RLUs/sec at λ_450_ divided by the RLUs/sec at λ_520_.

For testing the PAN^nita^ sensors in the SpLuc assay (**Figure 4C and Figure S16)**, a cosolvent of 42% methanol and 58% DMSO was used for all ligand dilutions except for heroin, which was dissolved solely in acetonitrile.

#### Titrations via yeast surface display

Ligand-specific titrations of nitazene sensors were carried out through yeast surface display, run similarly to library sorting, but with the following differences. A total of 10 µL of OD_600_=2 were labeled with various dilutions of ligand such that the ligand’s solvent was <1% of the reaction volume and ΔN-HAB1^T+^ was added to 500 nM in the final reaction volume of 50 µL. Data from 25,000 labeled cells was collected after resuspending cell pellets in 100 µL CBSF, processed in FlowJo v10 software, and fitted to a specific binding curve with Hill slope in GraphPad Prism 10 to find EC_50_ values.

---

**Fig. S1.**
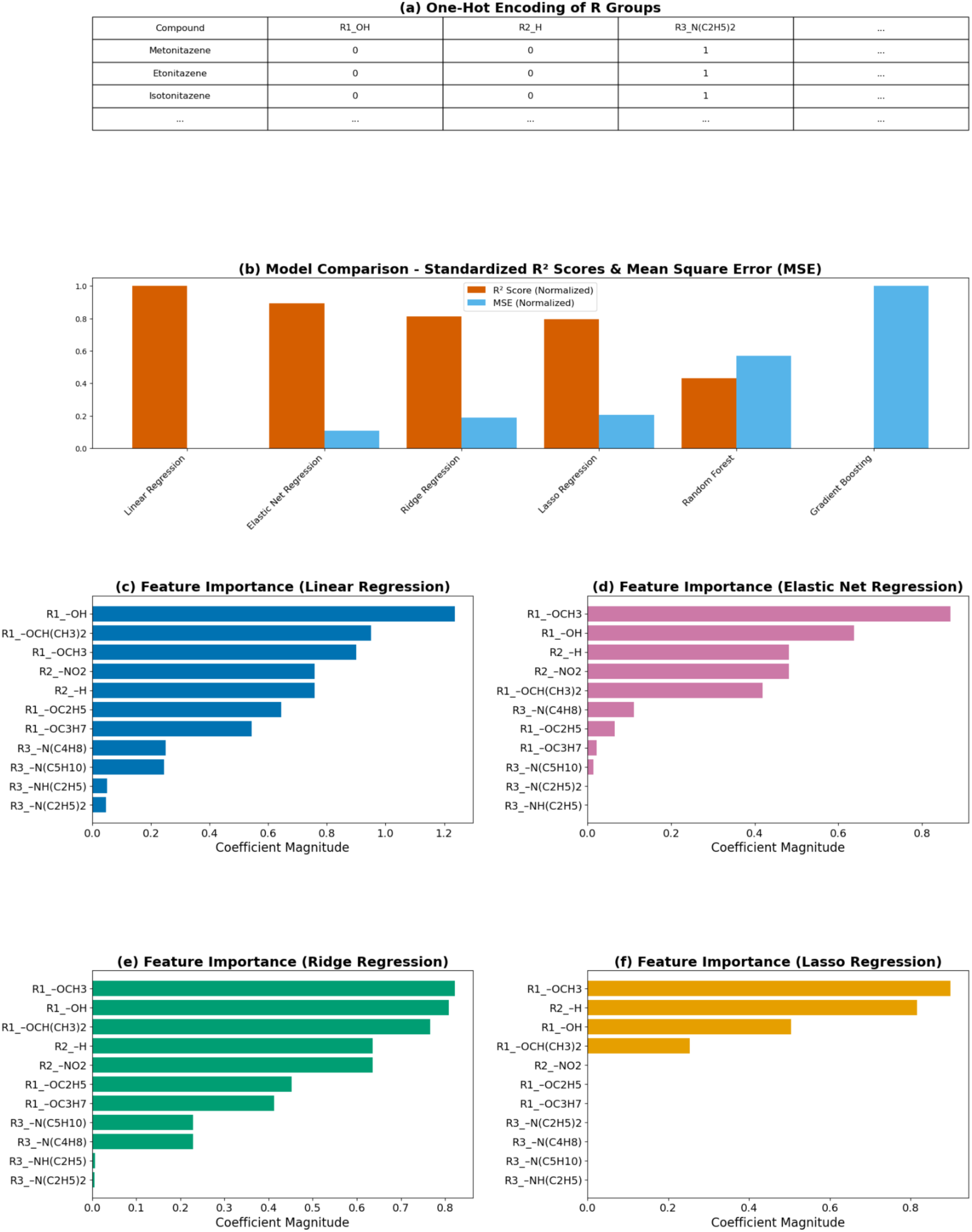
Summary of supervised learning of nitazene structural features. (a) One-hot encoding representation of R groups used as features in the regression models. (b) Model comparison showing standardized 𝑅^2^ scores (orange) and mean squared error (MSE, blue) for various regression models, with higher 𝑅^2^ and lower MSE indicating better model performance. (c-f) Feature importance for the top four performing models: (c) ordinary least squares linear regression, (d) elastic net regression, (e) ridge regression, and (f) lasso regression. The absolute magnitude of the learned regression coefficients indicate the relative contribution of each R group feature to the predicted 𝑙𝑜𝑔𝐸𝐶50 value (described here as the coefficient magnitude).

**Fig. S2.**
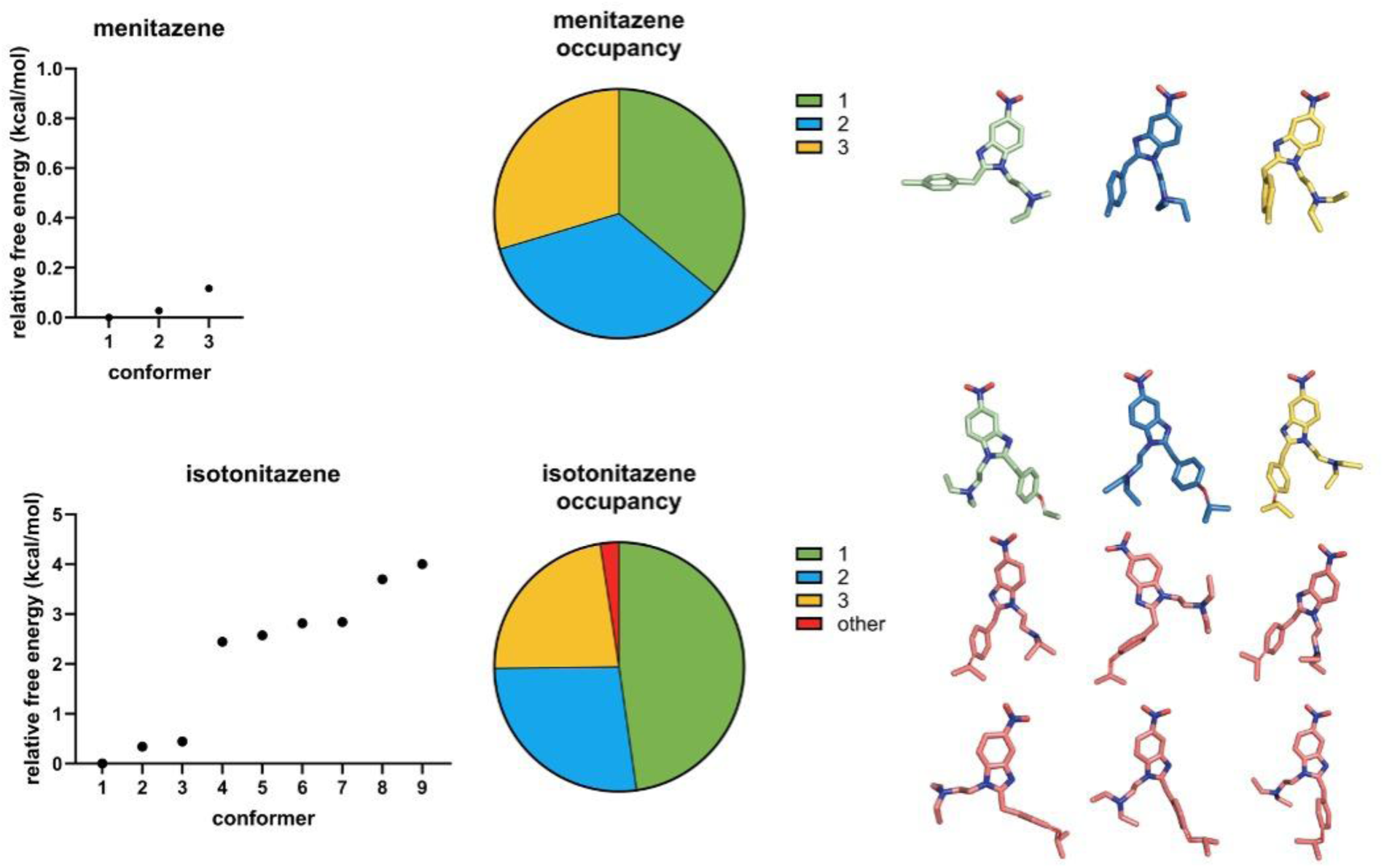
Menitazene and isotonitazene conformer selection by relative free energy and conformer occupancy.

**Fig. S3.**
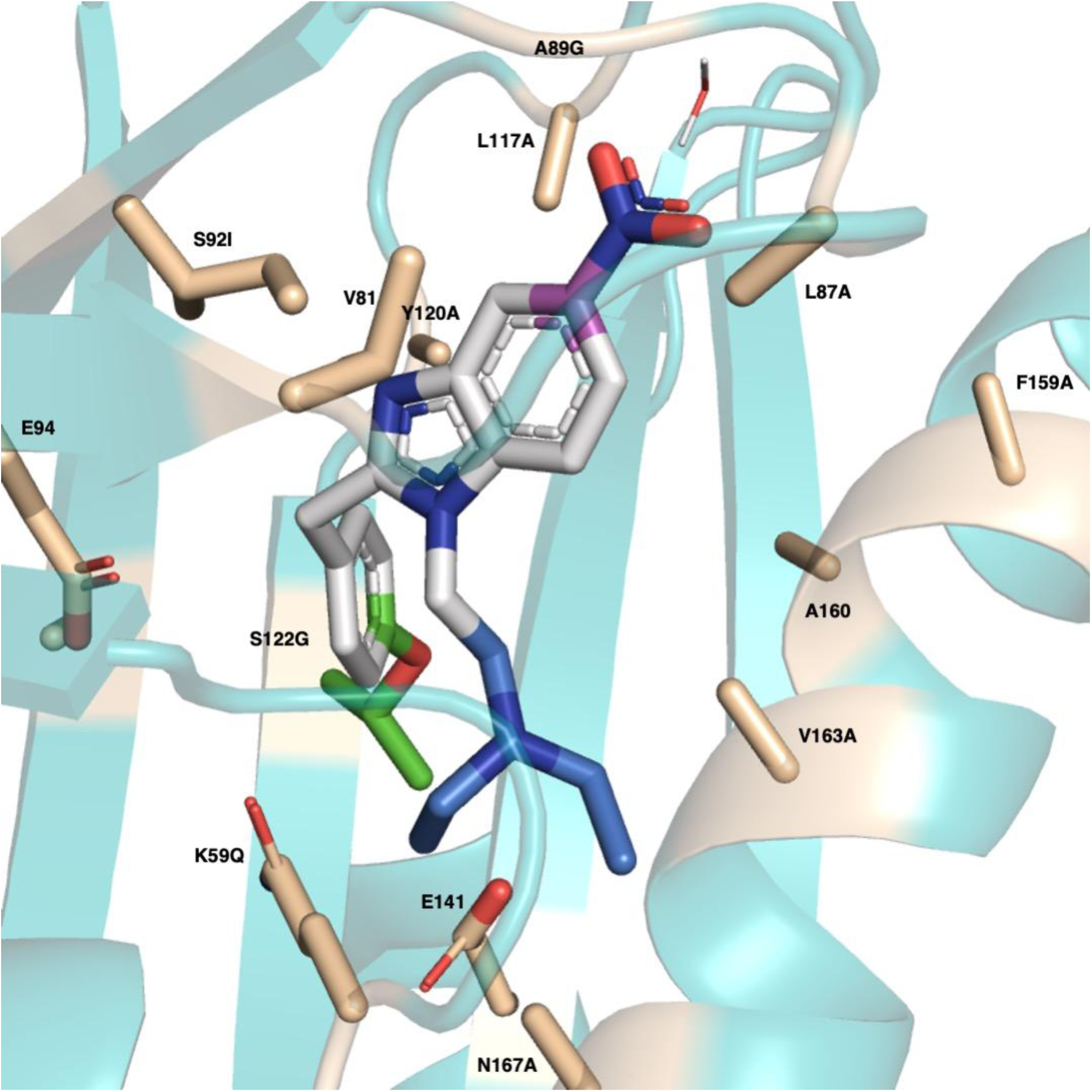
Isotonitazene conformers docked to the HMH library member with the largest allowable pocket volume (smallest amino acid contained in library at each position).

**Fig. S4.**
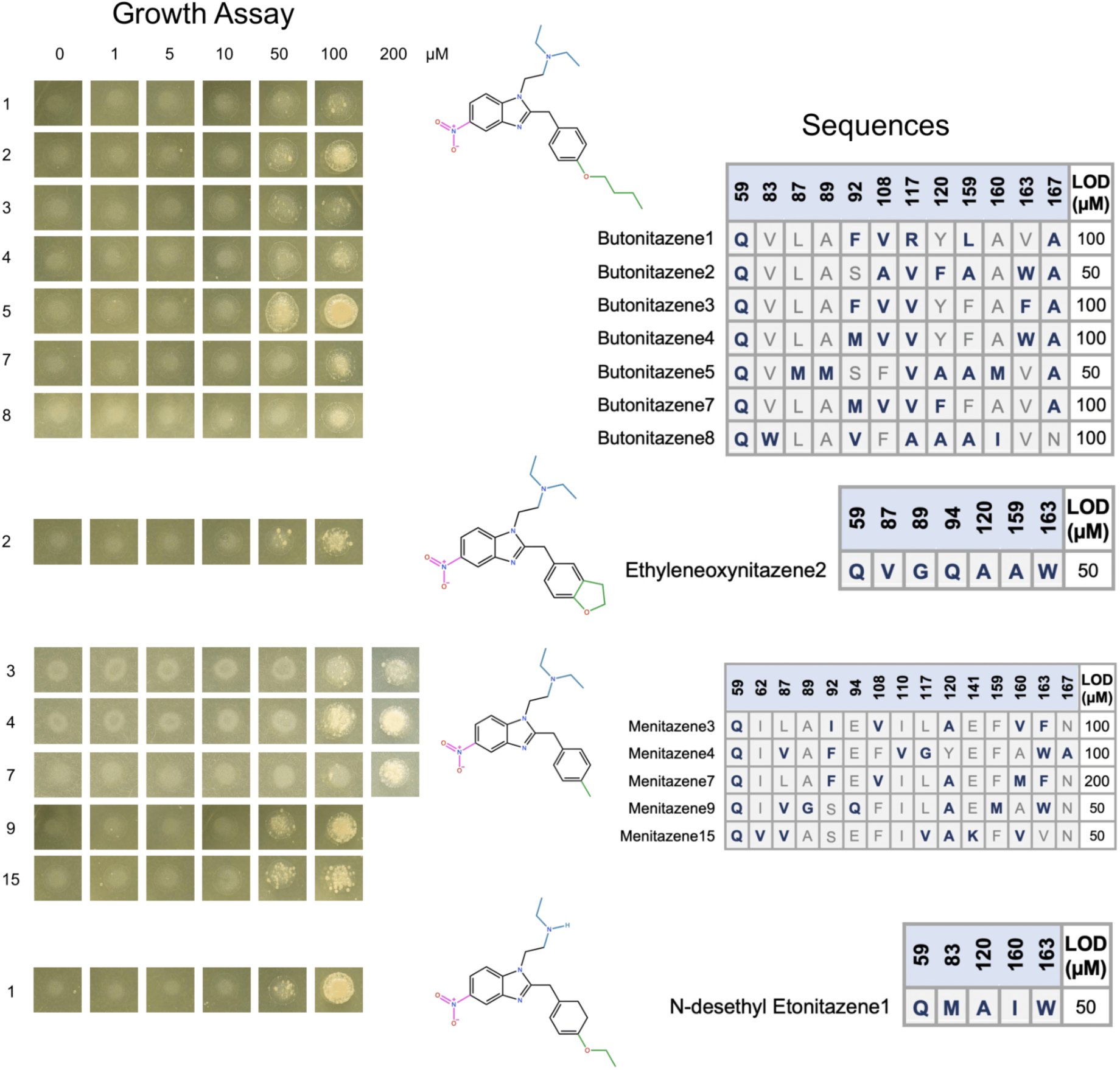
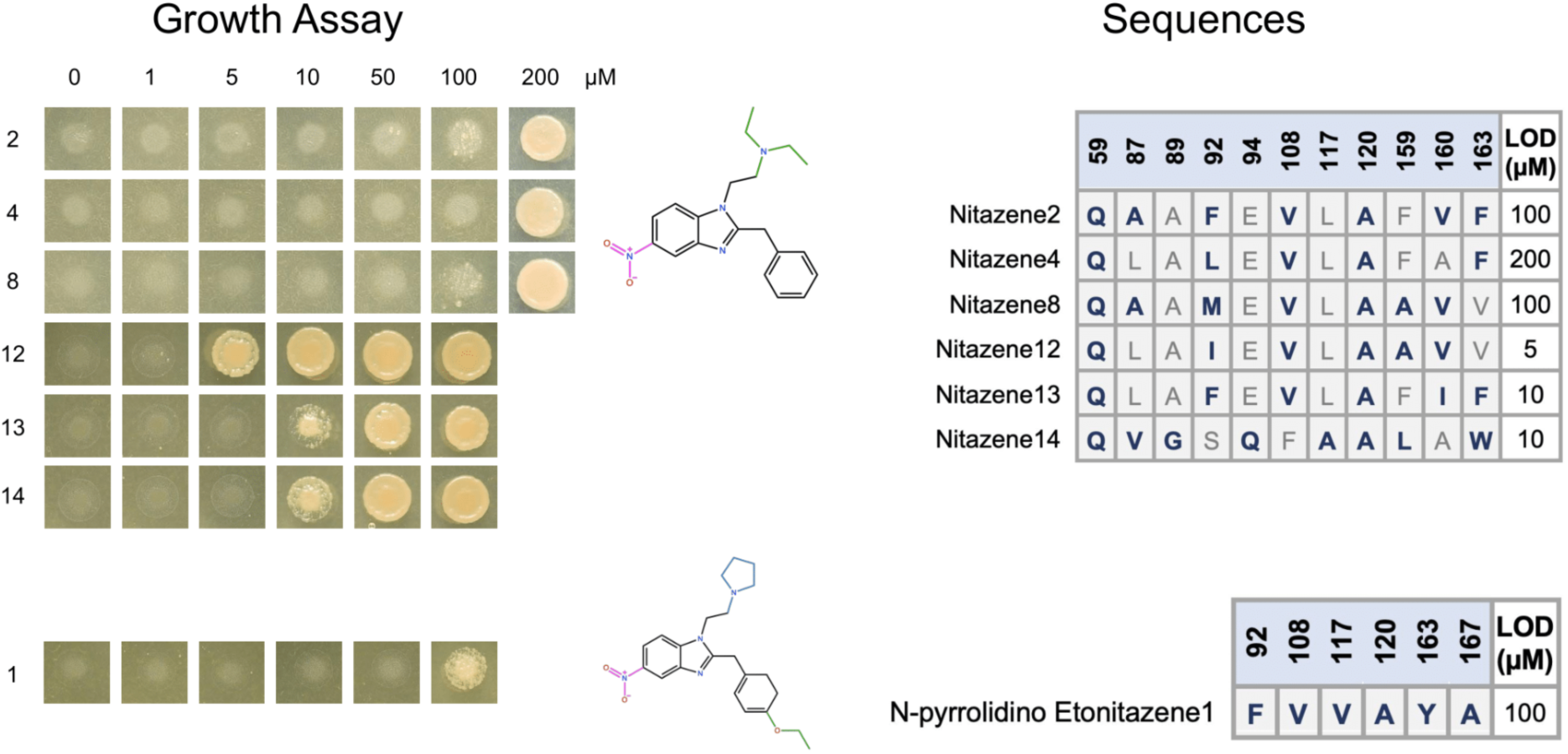
Initial Y2H nitazene hits from HMH library. Structure of nitazenes with growth assay of *S. cerevisiae* spots containing the respective biosensor and reporter circuit (MaV99 Y2H strain). Strains containing the chosen biosensor were spotted as 2 μL drops on SD -trp -leu -ura +ligand plates and allowed to grow for three days at 30 °C, then imaged. Growth indicates activation of the biosensor by the displayed nitazene. Amino acid substitutions for all biosensors is also shown. Blue letters indicate an amino acid substitution. Black numbers in the top row indicate the amino acid position in the PYR1 protein sequence. Library is in the PYR1 HOT5 background.

**Fig. S5.**
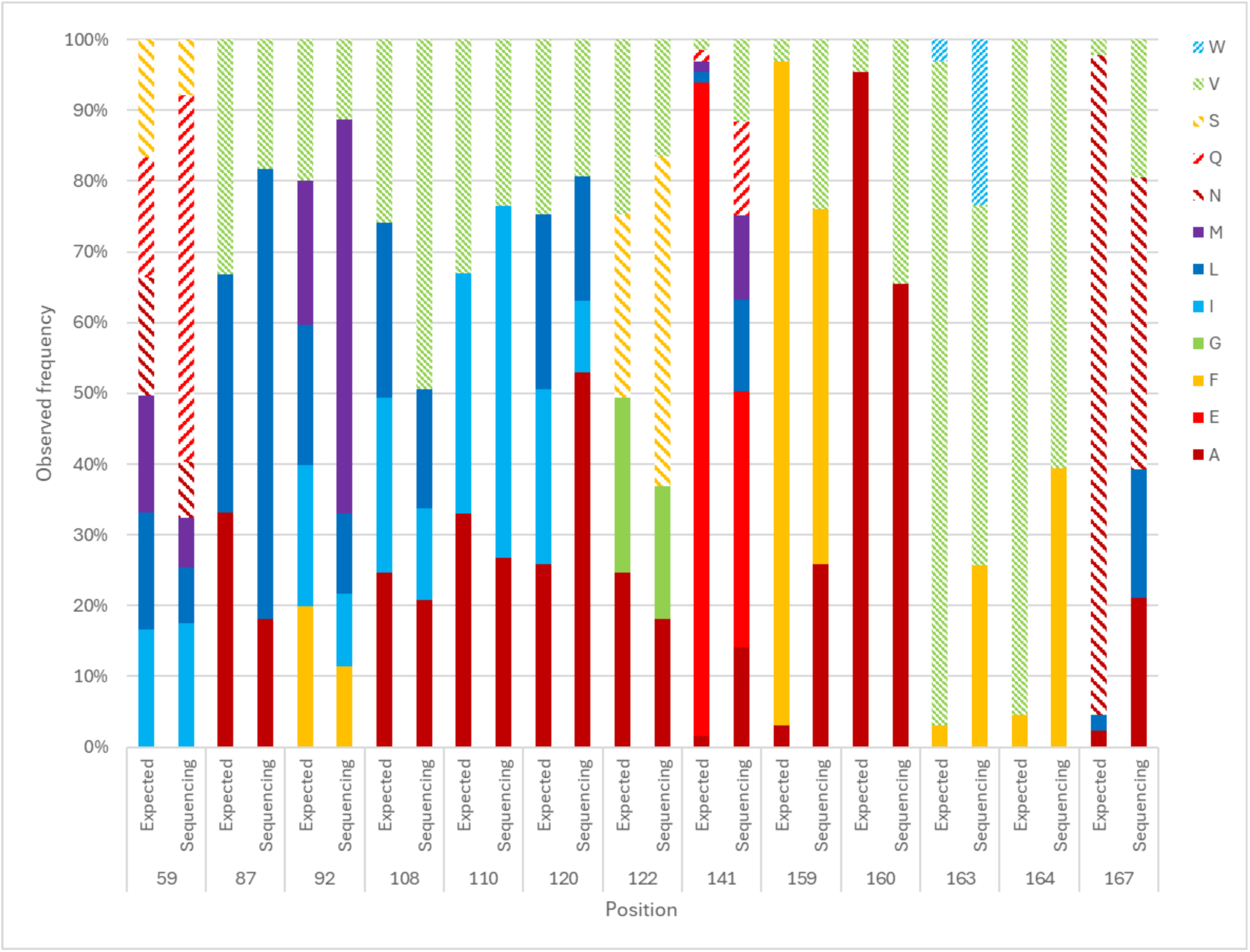
Amino Acid Distribution of NitazeneLib1. Design and sequencing results showing composition of NitazeneLib1. Plots compare the expected library amino acid distribution to the observed frequency calculated from library deep sequencing.

**Fig. S6.**
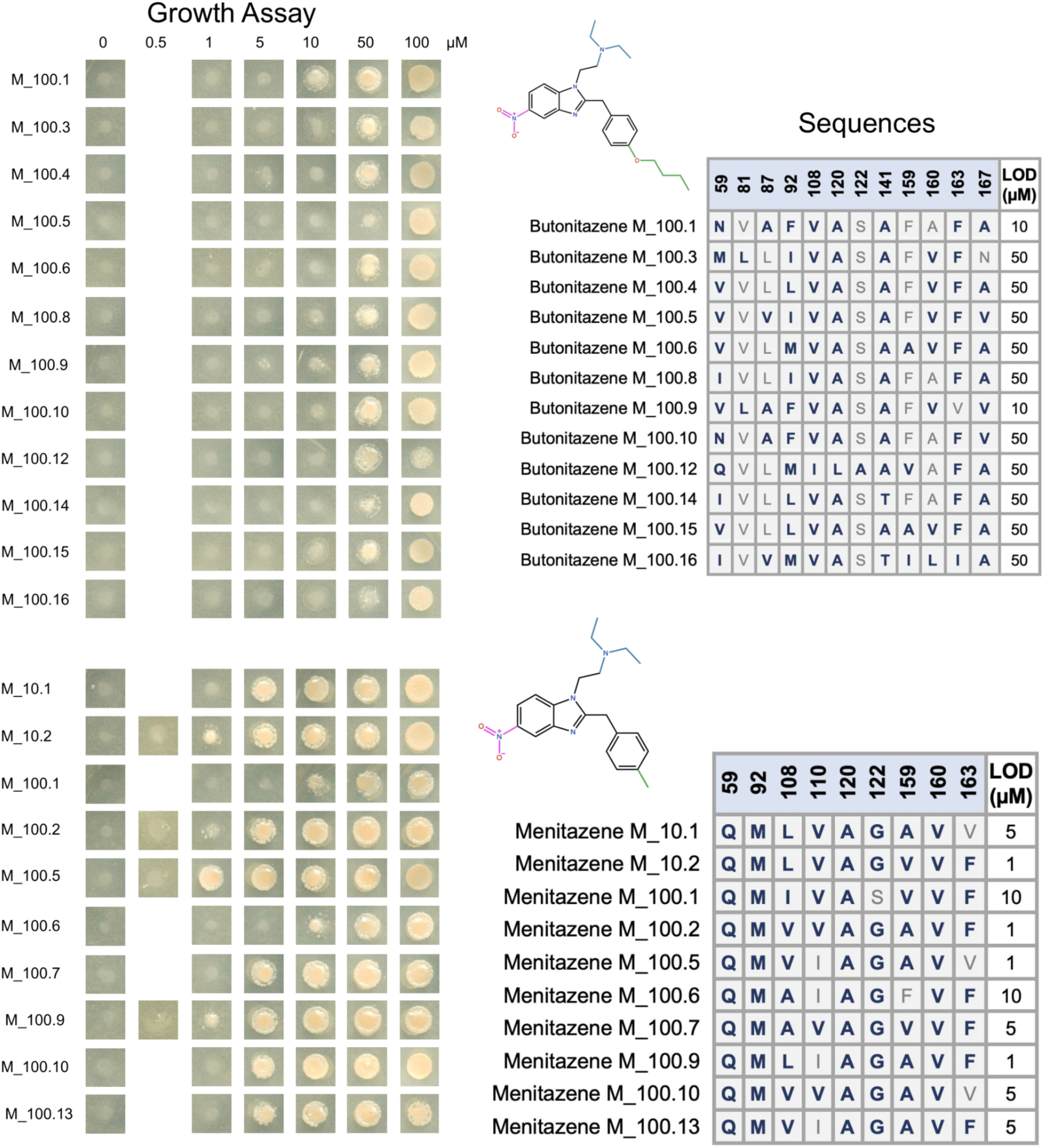
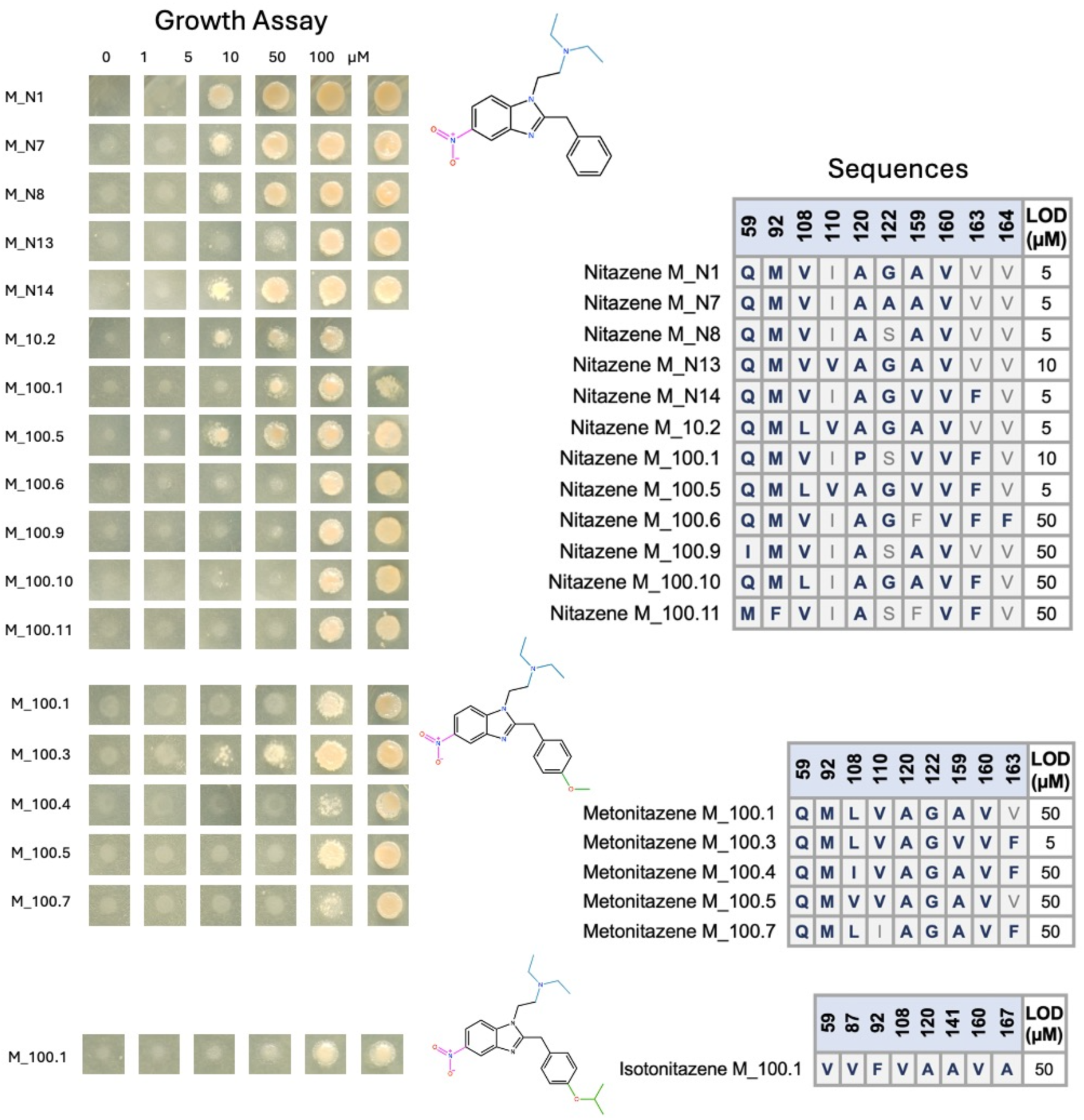
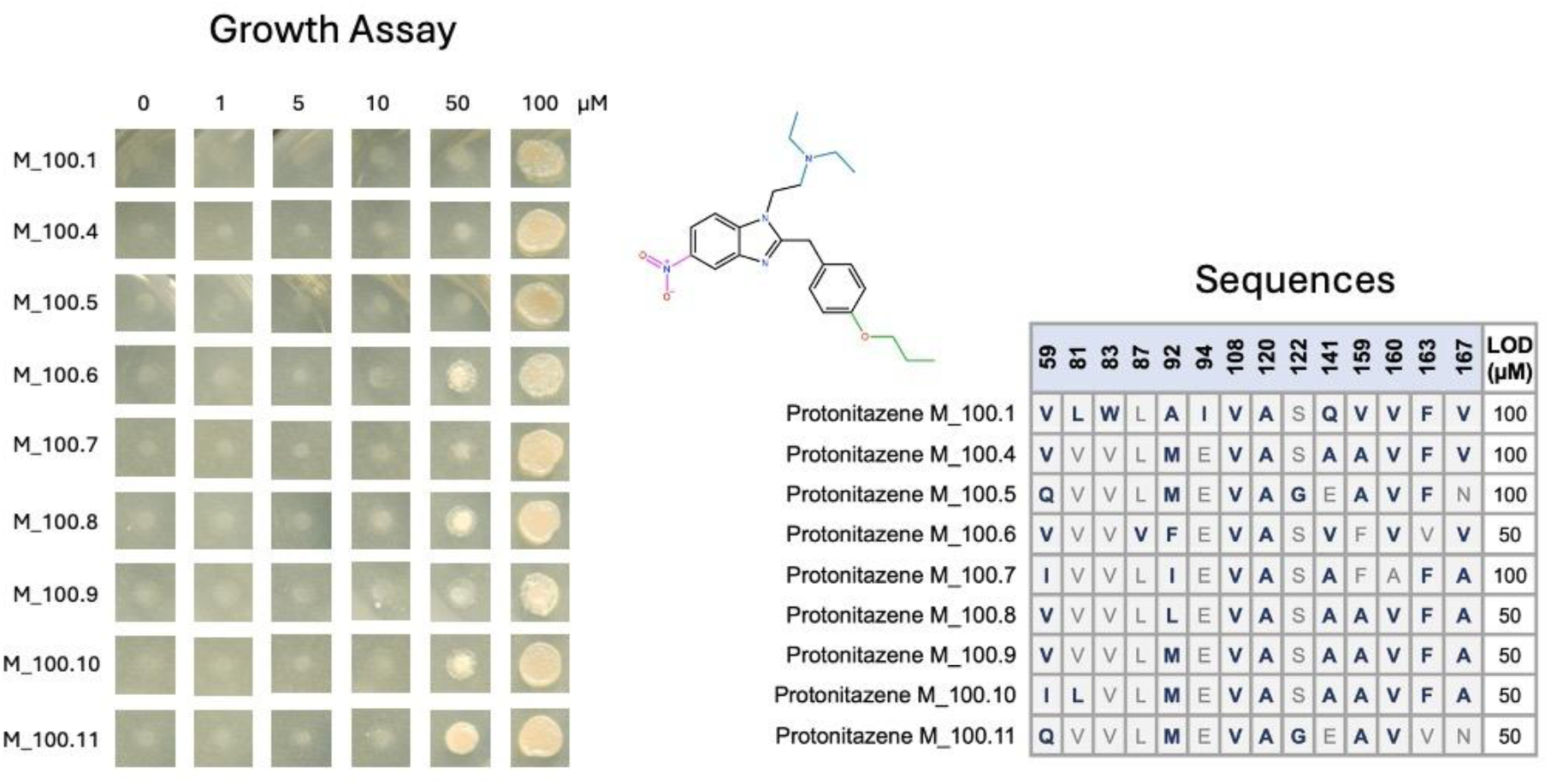
Complete Y2H results from NitazeneLib1. Structure of nitazenes with growth assay of S. cerevisiae spots containing the respective biosensor and reporter circuit (MaV99 Y2H strain). Strains containing the chosen biosensor were spotted as 2 μL drops on SD -trp -leu -ura +ligand plates and allowed to grow for three days at 30 °C, then imaged. Growth indicates activation of the biosensor by the displayed nitazene. Amino acid substitutions for all biosensors is also shown. Blue letters indicate an amino acid substitution. Black numbers in the top row indicate the amino acid position in the PYR1 protein sequence. The library is in the PYR1 HOT5 background (*14*).

**Fig. S7.**
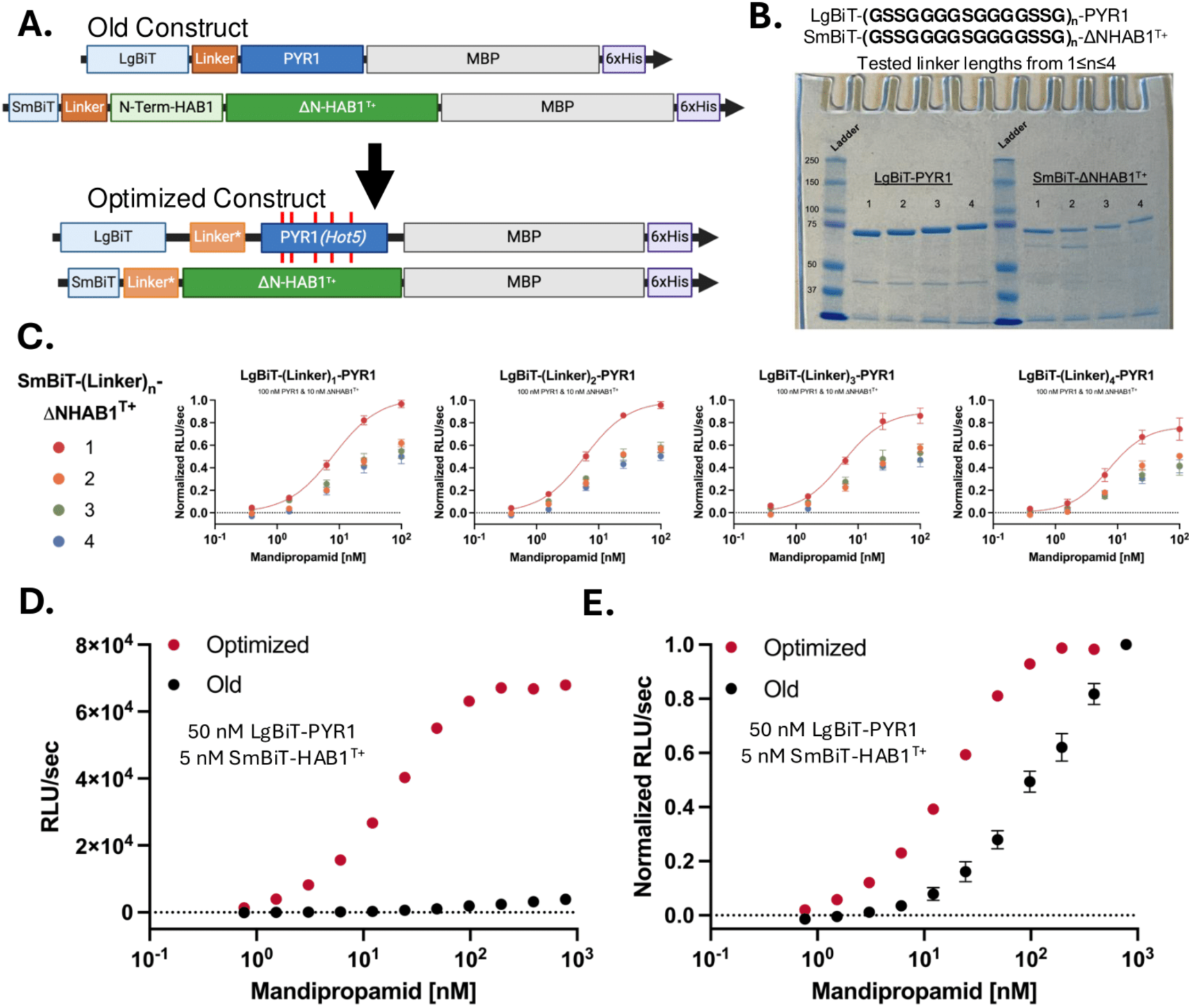
Optimization of LgBiT-PYR1 and SmBiT-HAB1 constructs for sensitive and specific light output. We added maltose binding protein domains to both constructs, varied the molar ratios of LgBiT-PYR1^nita^ to SmBiT-HAB1^T+^, tested a range of synthetic linker lengths connecting the split luciferase to its CID partner, and used a brighter luciferin. **A.** The SmBiT-HAB1 and LgBiT-PYR1 constructs were optimized for *E. Coli* expression and purification. PYR1 sensors were stabilized by including HOT5 mutations (Daffern et al. *J. Mol. Biol.* 2024)). The structurally disordered N-terminal region of HAB1 was removed to prevent proteolytic cleavage. 1-4 repeats of a 15 amino acid Gly/Ser rich linker (GS_2_G_4_SG_4_S_2_G)were encoded between LgBiT or SmBiT and PYR1 or HAB1 respectively. **B.** Each construct was purified via immobilized metal affinity chromatography, with purity visualized by SDS-PAGE and SimplyBlue^TM^ staining. **C.** All combinations of linker lengths were tested against one another. The shorter linkers tend to exhibit higher signal, with this trend being most pronounced between SmBiT-ΔNHAB1^T+^ variants. We selected the shortest linker (15 AA, n=1) for future experiments for both LgBiT-PYR1 and SmBiT-ΔNHAB1^T+^ and fitted those data to a curve here. **D.** The optimized constructs offer over 20-fold higher signal than the old constructs. **E.** The optimized constructs have higher affinity for one another shown here by decreased EC_50_. All data points represent the mean of replicate data (Fig. C–n = 3, Fig. D,E–n=4) and error bars represent the standard error of the mean.

**Fig. S8.**
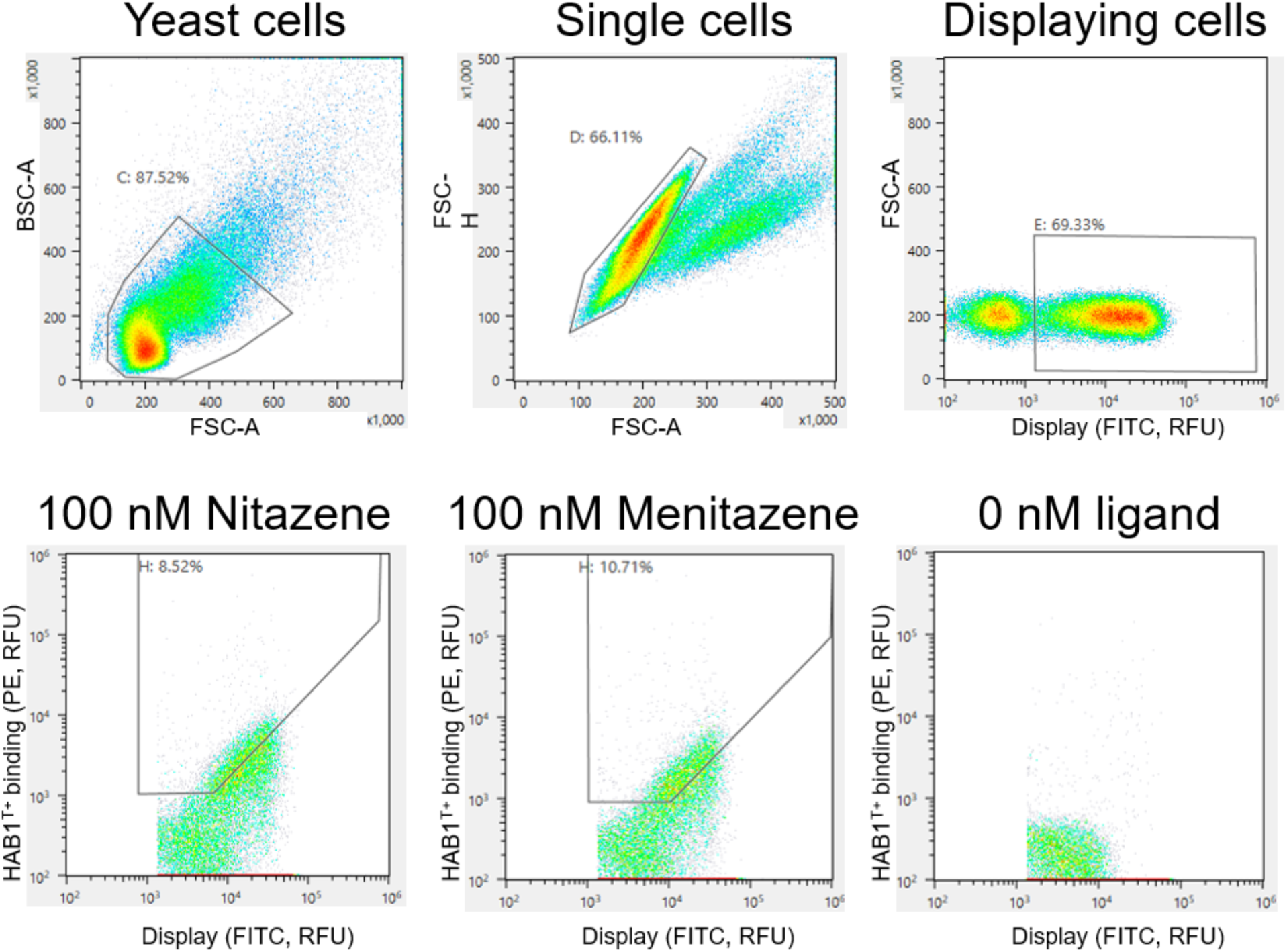
Example sorting gates for deep mutational scanning of PYR1^nita^. The top three cytograms include gates that were consistent for all sorts and the bottom three cytograms include gates used for sorting the three conditions for deep mutational scanning.

**Fig. S9.**
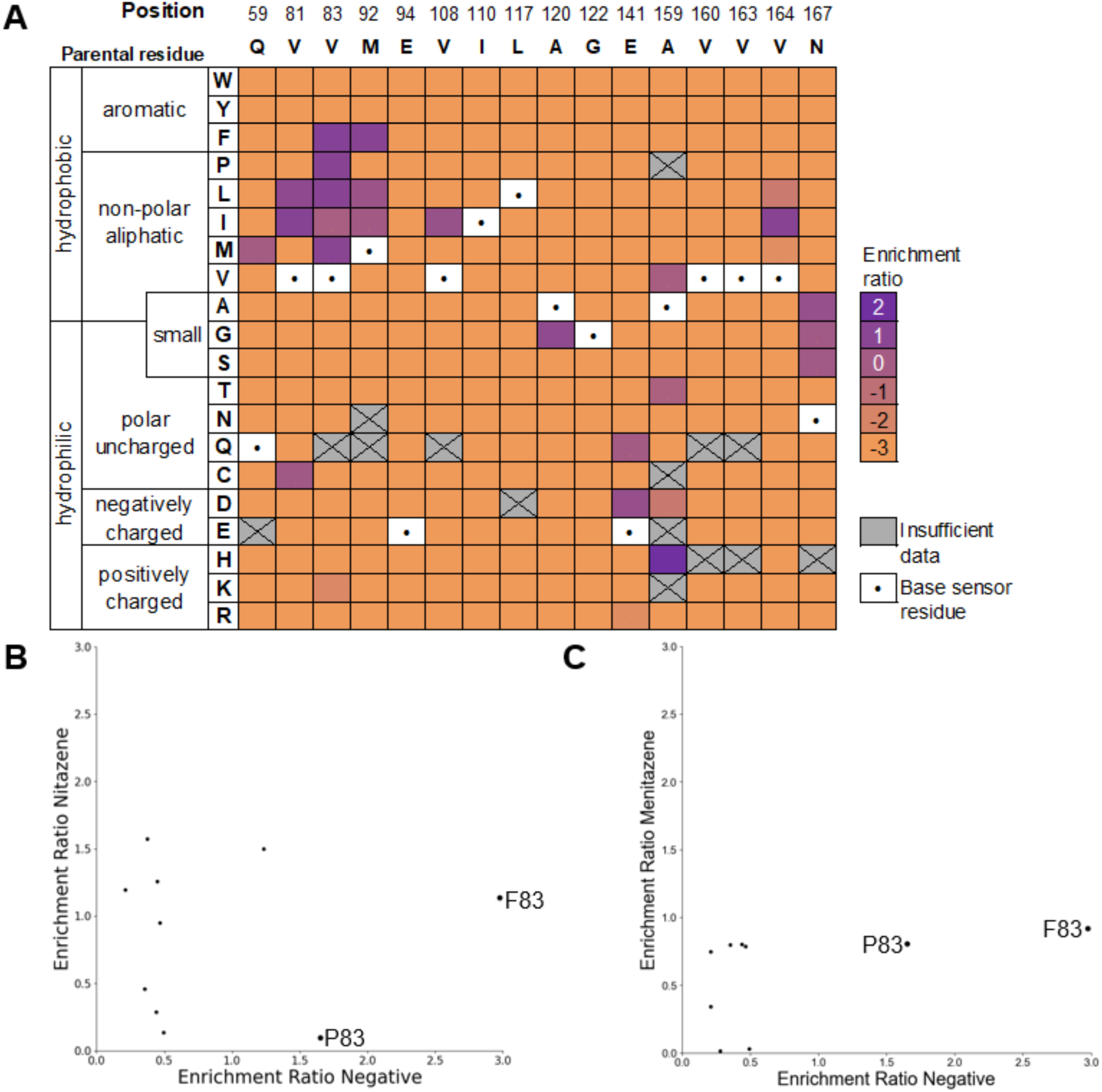
Deep mutational scans of PYR1nita on menitazene and no ligand control (negative). **A**. Heatmap of per-position normalized enrichment ratios for binding to menitazene. **B,C**. No ligand binding sort compared to (B) nitazene and (C) menitazene sort enrichment ratios. The no ligand binding (negative) occurred with 100 nM biotinylated ΔN-HAB1^T+^ and 1% (v/v) DMSO. This negative sort enriches mutations that bind *ΔN-HAB1^T+^* in the absence of ligand.

**Fig S10.**
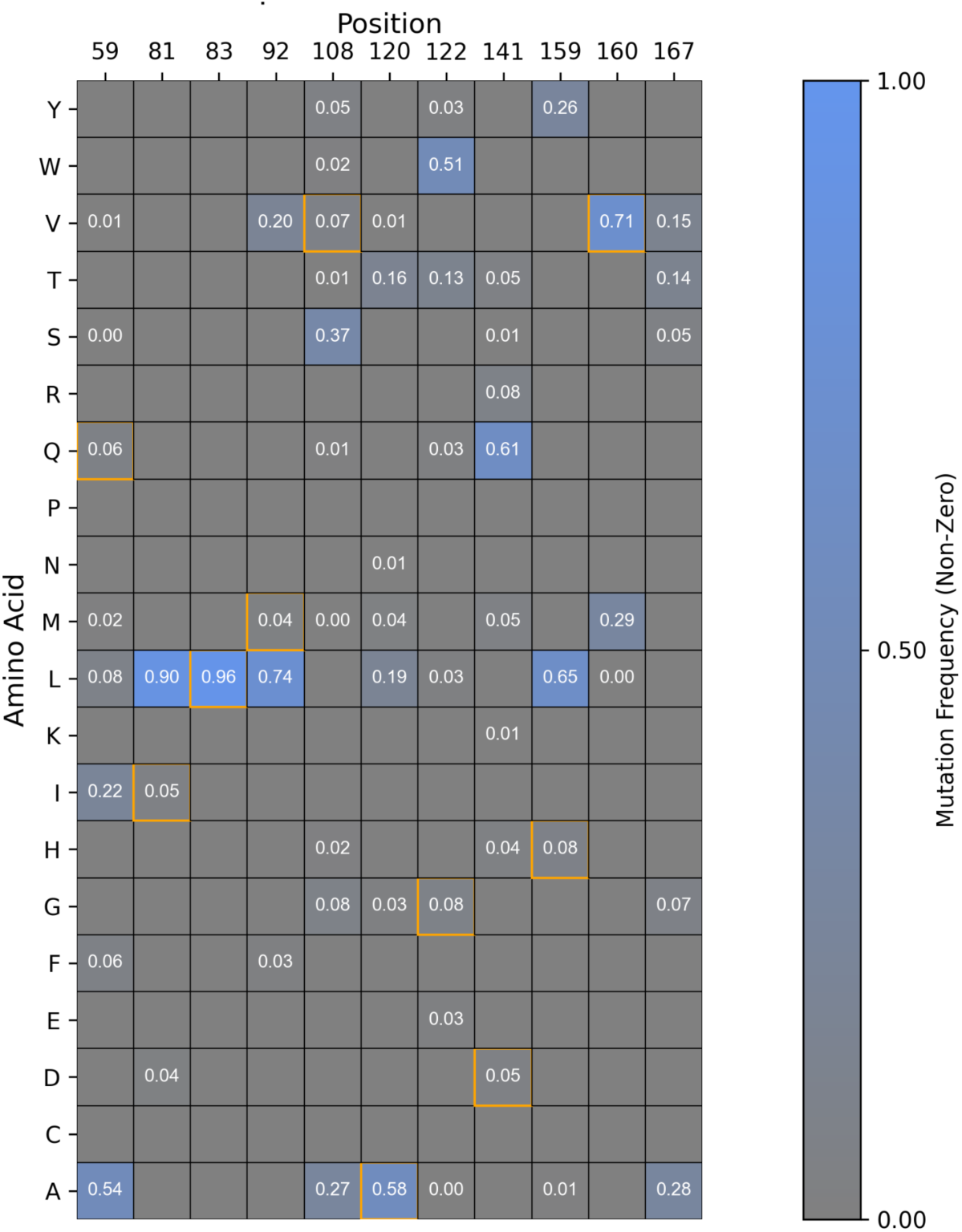
Heatmap of mutation frequencies at selected positions after LigandMPNN redesign. 224 redesigns of the PYR1 pocket for binding to nitazene. White numbers correspond to all non-zero frequencies for a given mutation. Orange boxes indicate the mutations present in the PYR^nitav2^ sensor.

**Fig. S11.**
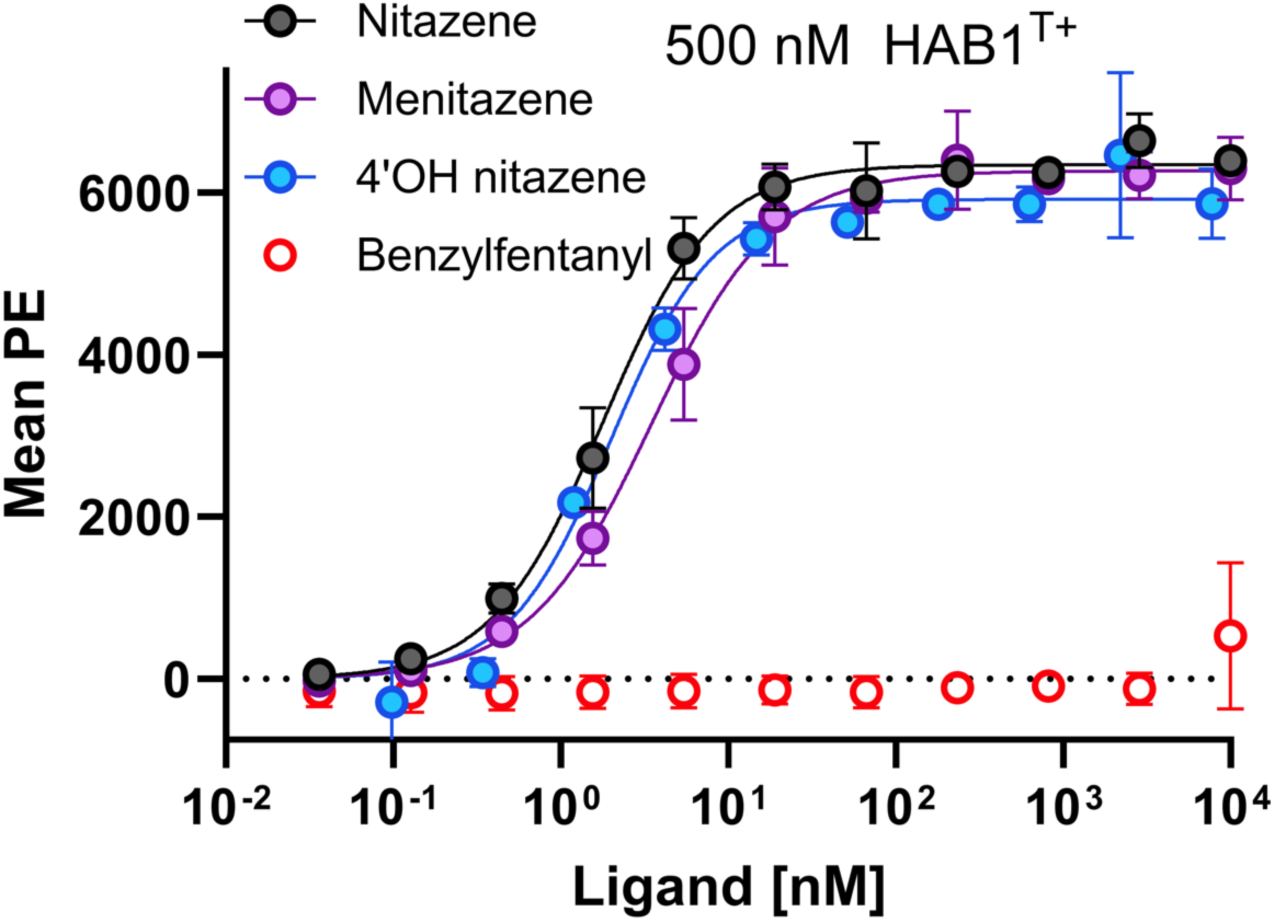
Yeast surface display titration of 4-hydroxy nitazene titration. Sensor PYR1^nitav2.1^ was titrated against 12 concentrations of nitazene, menitazene, 4-hydroxy nitazene, and benzylfentanyl with 500 nM biotinylated ΔN-HAB1^T+^. The EC_50_ of this sensor for each of the ligands is as follows: 1.8 nM nitazene, 3.5 nM menitazene, 2.0 nM 4-hydroxy nitazene. Error bars represent 1 s.d. (n=4).

**Fig. S12.**
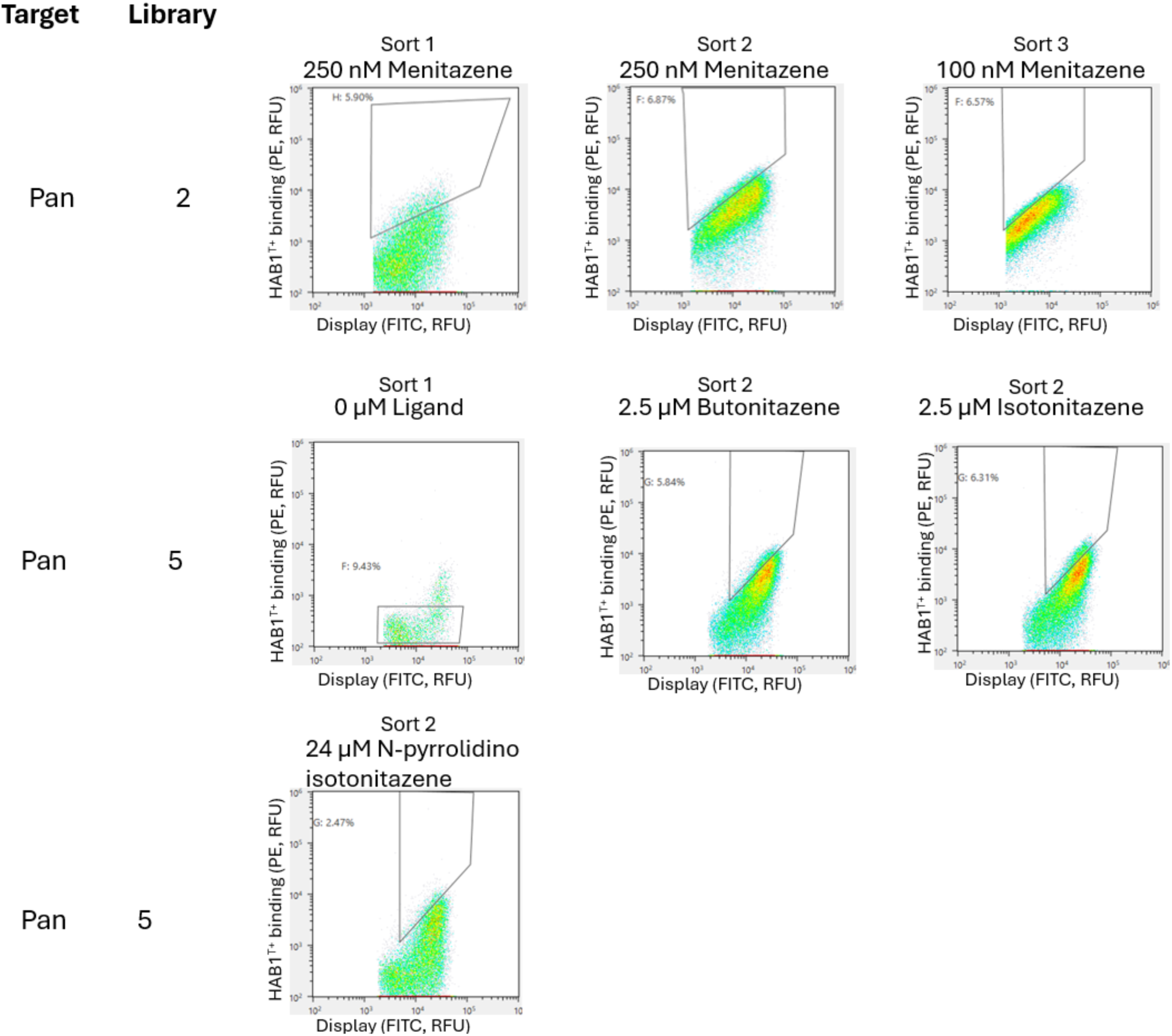
Sorting gates for acquiring PAN^nita^ sensors. Nitazene library 2 was sorted against menitazene, 3 times with 250 nM twice and once with 100 nM. Nitazene library 5 was sorted first with no ligand at 1% DMSO to collect non-constitutive binders, then sorted in parallel with 2.5 uM of both butonitazene and isotonitazene and 24 uM N-pyrrolidino isotonitazene. 200,000 cells were collected except for 24 uM N-pyrrolidino isotonitazene where 164,000 cells were collected due to limited numbers of cells above background.

**Fig. S13.**
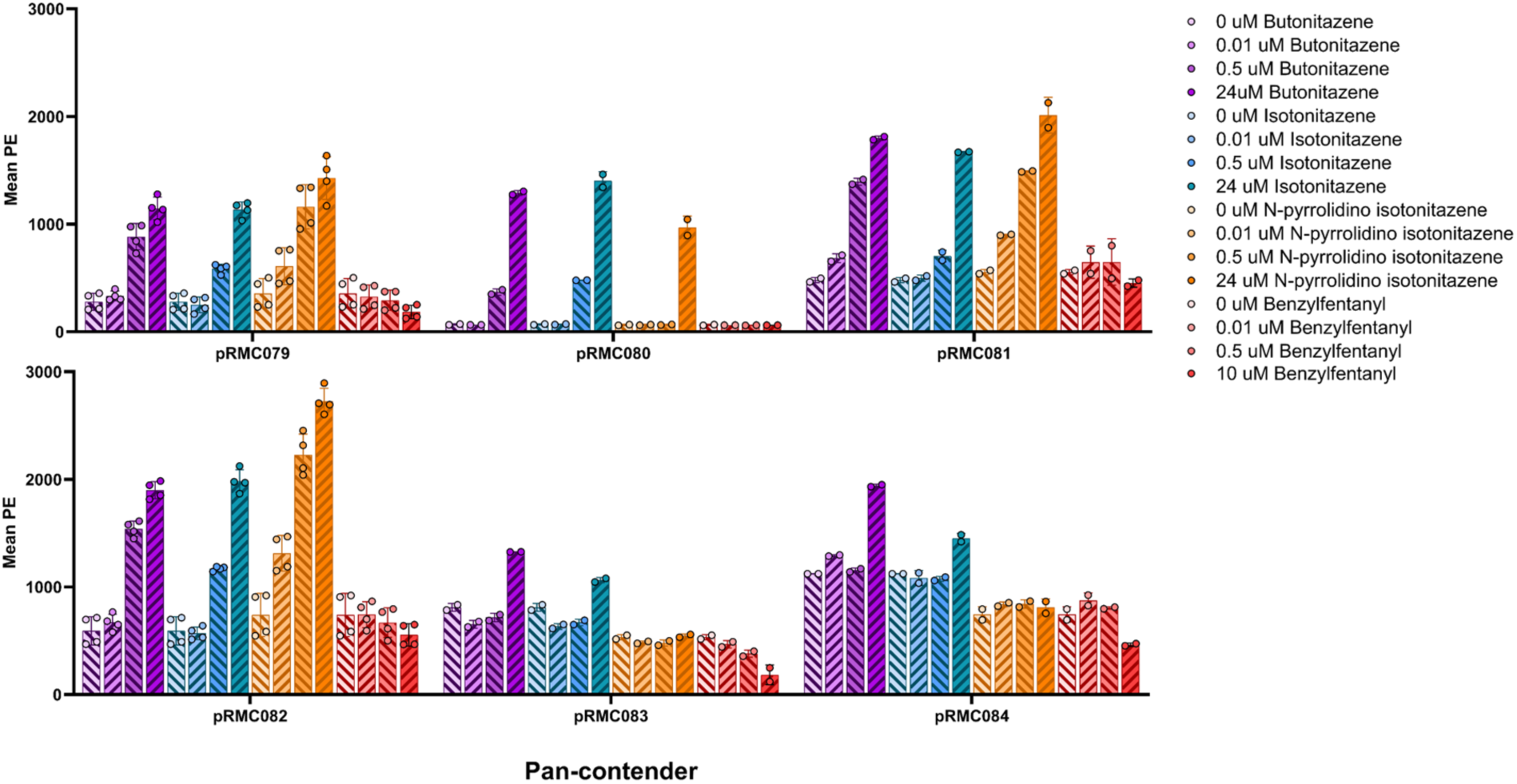
Yeast surface display binding for all pan-nitazene contenders measured for indicated ligands. pRMC079 - pRMC084 were screened against the indicated concentrations of butonitazene, isotonitazene, N-pyrrolidino isotonitazene, and benzylfentanyl. pRMC079 is PAN^nita^ and pRMC082 is PAN^nita.1^, which were selected as the top contenders due to their either low constitutive activity or high dynamic range.

**Fig. S14.**
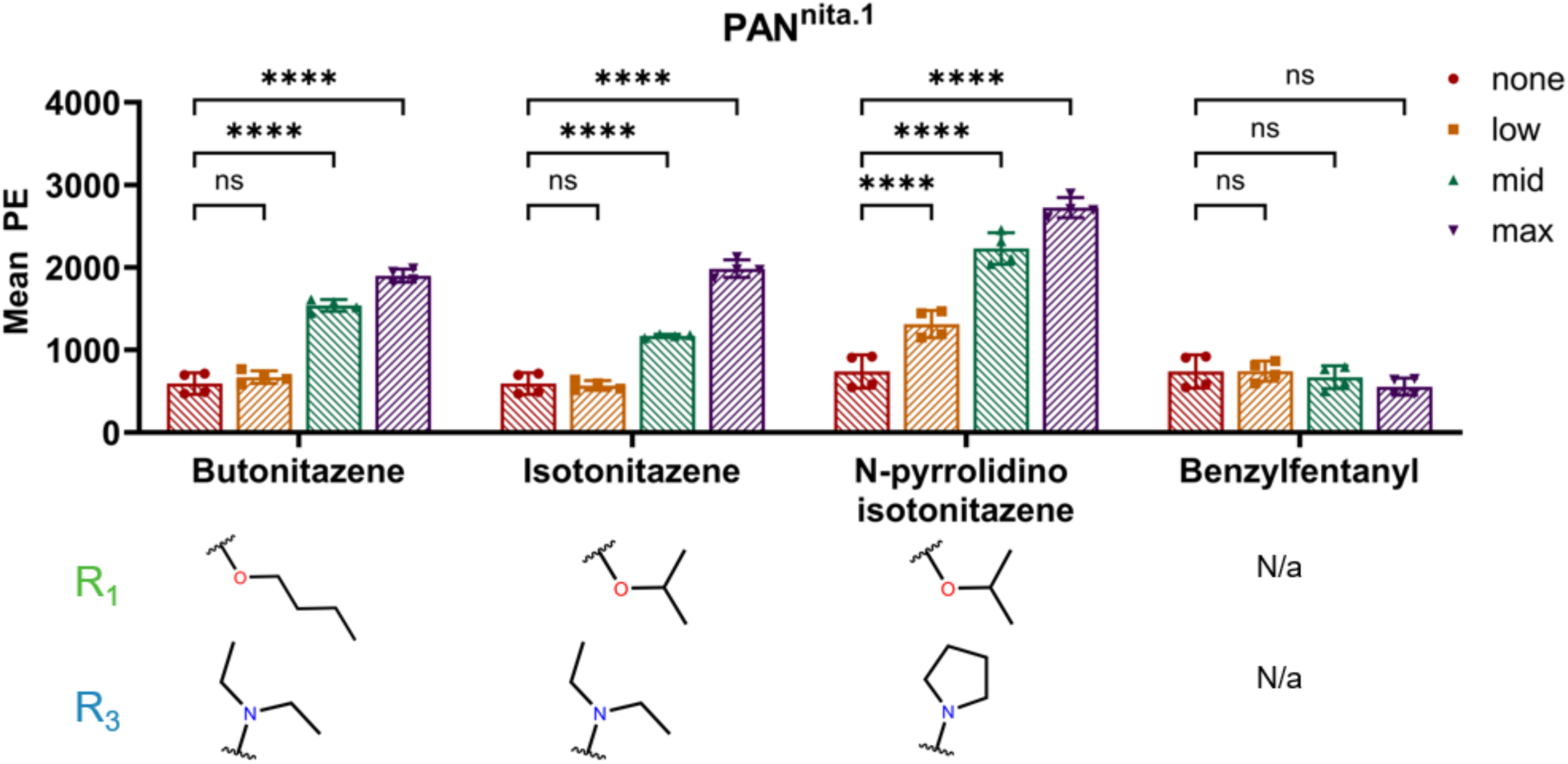
Yeast surface display binding for PAN^nita.1^ measured for indicated ligands. **** indicates p<0.0001 and ns indicates not significant. n=4, error bars indicate 1 s.d.

**Fig. S15.**
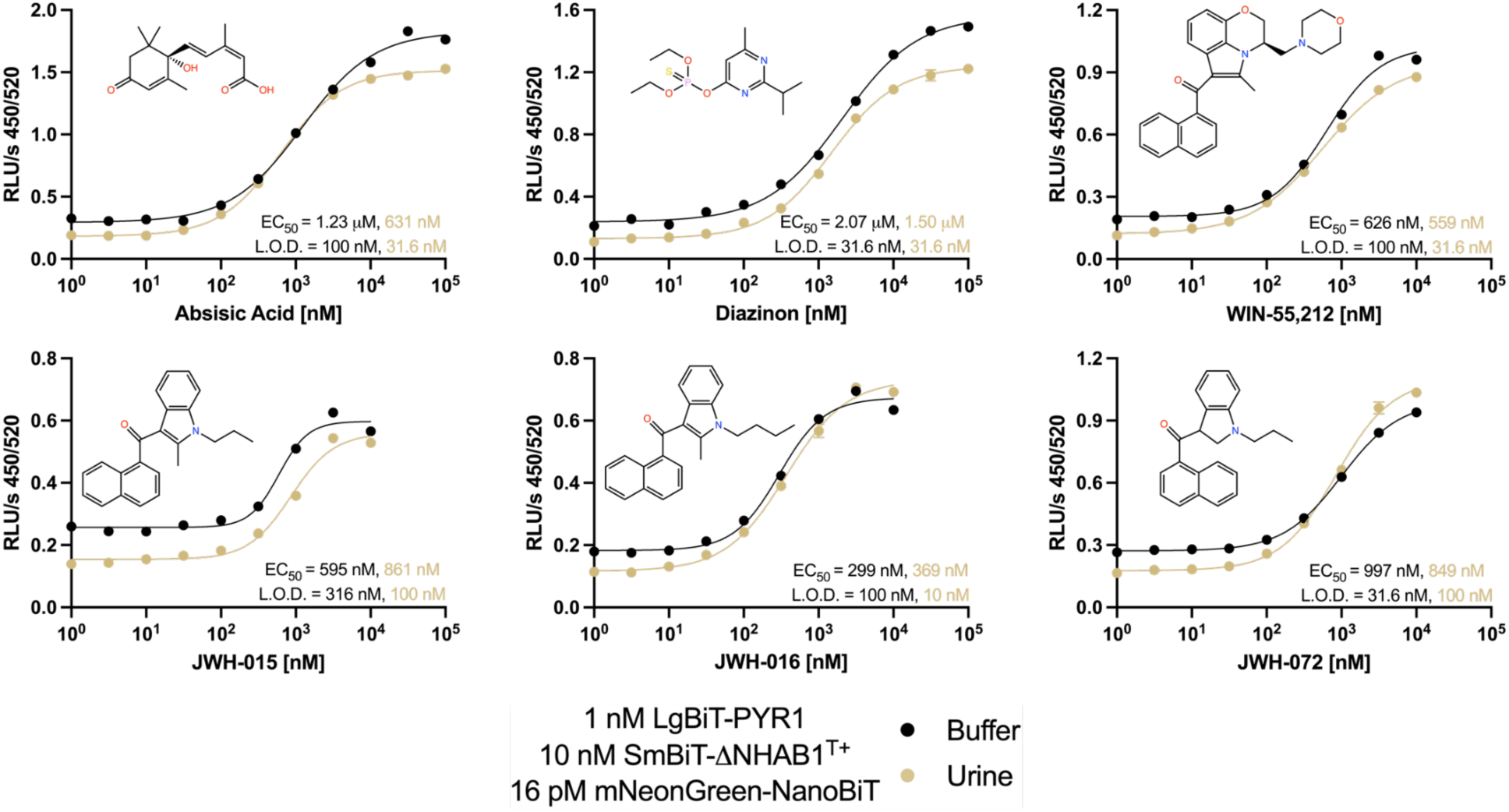
Validation of split luciferase diagnostic assay. Ratiometric luminescence assay for indicated ligands. Protein concentrations are indicated, and black (buffer) and gold (urine) circles represent data series. Solid lines represent best fit of a sigmoidal function (Hill coefficient =1). All measurements were collected in triplicates (n=3) and error bars are shown 1 s.e.m.. In all cases, error bars are smaller than the symbol.

**Fig. S16.**
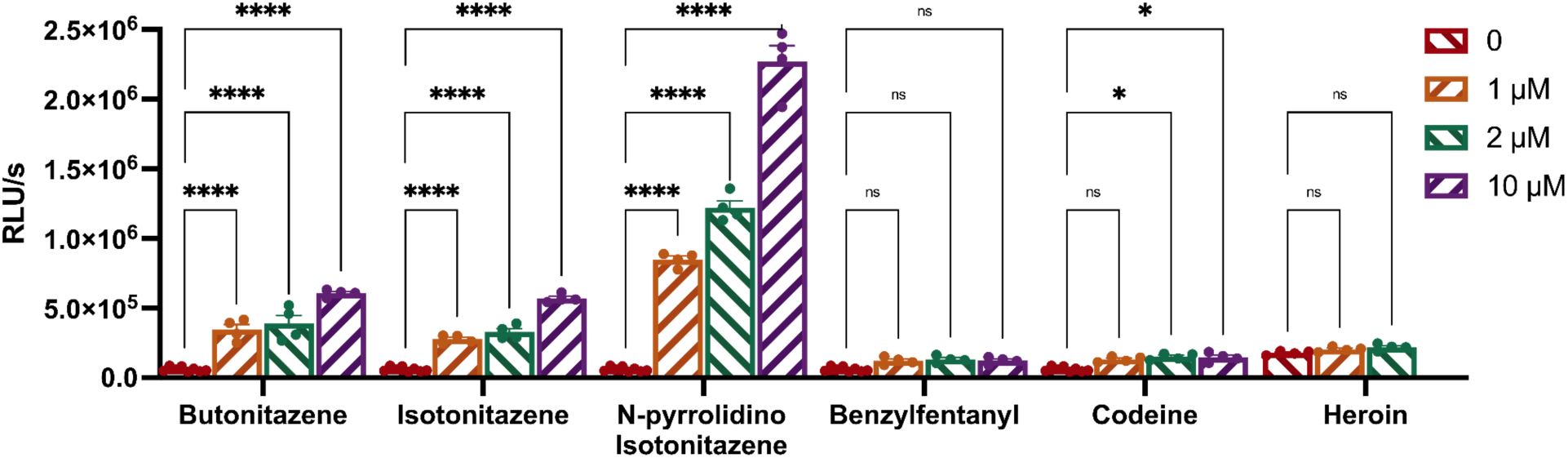
Split luciferase assay data for PAN^nita^ measured for indicated ligands. We utilize 4 nM LgBiT-PYR1 and 4 nM SmBiT-ΔNHAB1. **** p<0.0001, * p<0.05, ns not statistically significant. Error bars are shown for 1 S.E.M. for n=4 replicates.

**Table S1.**
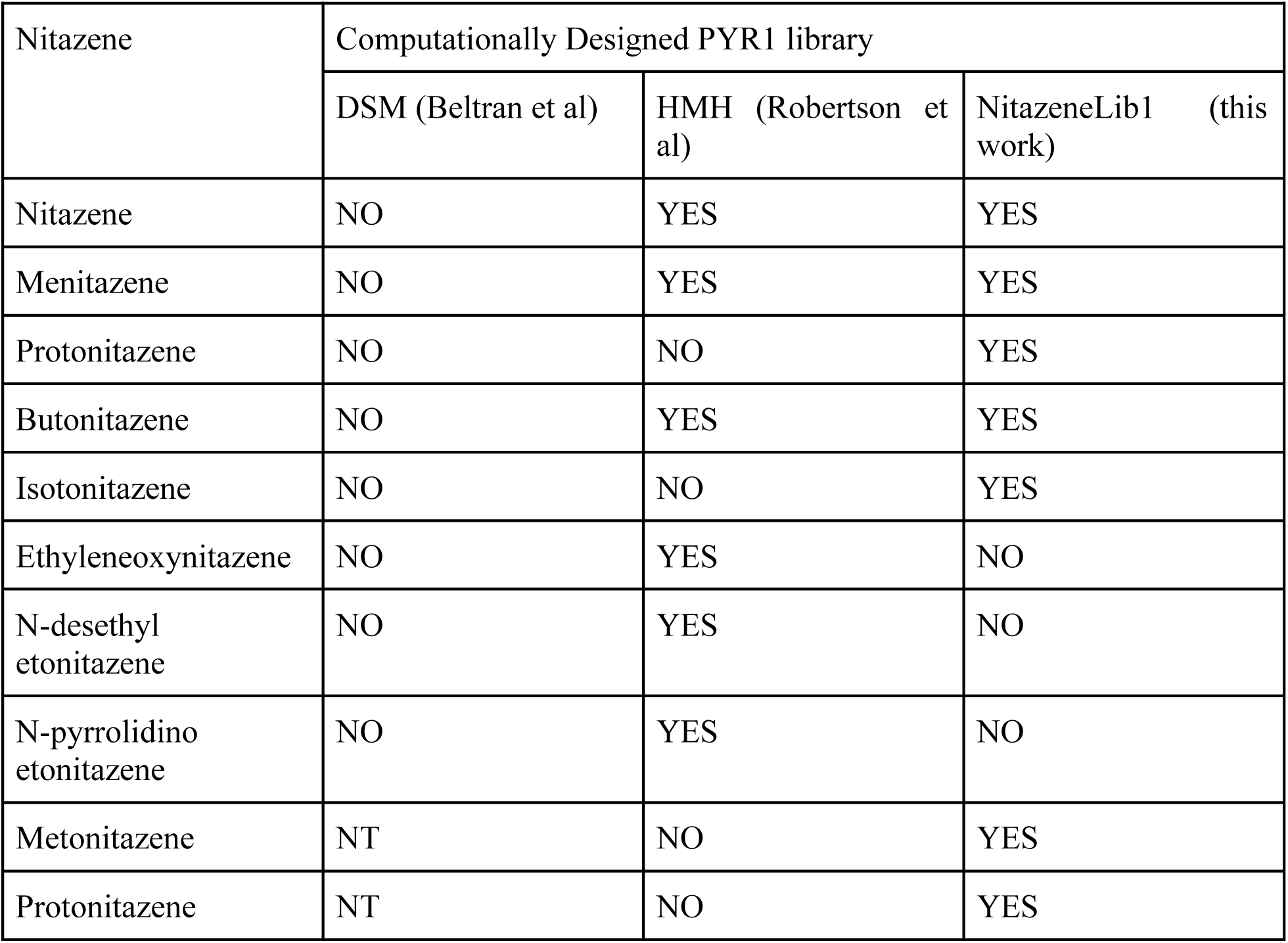
Panel of nitazenes screened in yeast two hybrid selections. YES indicates at least one sensor sequence was responsive to the indicated nitazene at 100 𝜇M, NO indicates that no sequence was found, NT indicates that the condition was not tested.

**Table S2.** HMH library input and nitazene screen output.

| WT | K59 | V83 | L87 | A89 | S92 | E94 | F108 | I110 | L117 | Y120 | S122 | E141 | F159 | A160 | V163 | N167 |
| --- | --- | --- | --- | --- | --- | --- | --- | --- | --- | --- | --- | --- | --- | --- | --- | --- |
| HMH library | KQR | VW | ALV | AG | FIMS | EKQ | AFV | I | ALV | ARVY | GLS | EK | AFILMT | AIVM | AFMWY | AN |
| Butonitazene1 | Q |  |  |  | F |  | V |  | R |  |  |  | L |  |  | A |
| Butonitazene2 | Q |  |  |  |  |  | A |  | V | F |  |  | A |  | W | A |
| Butonitazene3 | Q |  |  |  | F |  | V |  | V |  |  |  |  |  | F | A |
| Butonitazene4 | Q |  |  |  | M |  | V |  | V |  |  |  |  |  | W | A |
| Butonitazene5 | Q |  | M | M |  |  |  |  | V | A |  |  | A | M |  | A |
| Butonitazene7 | Q |  |  |  | M |  | V |  | V | F |  |  |  |  |  | A |
| Butonitazene8 | Q | W |  |  | V |  |  |  | A | A |  |  | A | I |  |  |
| Ethyleneoxynitazene<br>2 | Q |  | V | G |  | Q |  |  |  | A |  |  | A |  | W |  |
| Menitazene3 | Q |  |  | I |  |  | V |  |  | A |  |  |  | V | F |  |
| Menitazene4 | Q |  | V |  | F |  |  | V | G |  |  |  |  | W | A |  |
| Menitazene7 | Q |  |  |  | F |  | V |  |  | A |  |  |  | M | F |  |
| Menitazene9 | Q |  | V | G |  | Q |  |  |  | A |  |  | M |  | W |  |
| Menitazene15 | Q |  | V | G |  | Q |  |  | A | A |  |  |  |  | W | A |
| N-desethyl<br>Etonitazene1 | Q | M |  |  |  |  |  |  |  | A |  |  |  | I | W |  |
| N-pyrrolidino<br>Etonitazene1 |  |  |  |  | F |  | V |  | V | A |  |  |  |  | Y | A |
| Nitazene2 | Q |  | A |  | F |  | V |  |  | A |  |  |  | V | F |  |
| Nitazene4 | Q |  |  |  | L |  | V |  |  | A |  |  |  |  | F |  |
| Nitazene8 | Q |  | A |  | M |  | V |  |  | A |  |  | A | V |  |  |
| Nitazene9 | Q |  | V | G |  |  |  | V |  | A |  |  | M |  | W |  |
| Nitazene12 | Q |  |  |  | I |  | V |  |  | A |  |  | A | V |  |  |
| Nitazene13 | Q |  |  |  | F |  | V |  |  | A |  |  |  | I | F |  |
| Nitazene14 | Q |  | V | G |  | Q |  |  | A | A |  |  | L |  | W |  |

**Table S3.** Benchmarking PYR1 ligands during structural replacement docking.

| Ligand | Mutations from WT<br>PYR1 | Total conformers<br>observed in<br>simulation | Conformer<br>energy<br>cutoff<br>(kcal/mol) | Conformers<br>under energy<br>cutoff | Alignments<br>per<br>conformer | Total<br>alignments<br>sampled | Alignments<br>fitting<br>sequence |
| --- | --- | --- | --- | --- | --- | --- | --- |
| mandipropamid | WT | 185 | 0.5 | 14 | 400 | 5600 | 0 |
|  | K59R V81I<br>F108A S122G<br>F159L |  |  |  | 80 | 1120 | 1 |
| WIN55,212-2 | WT | 51 | 0.5 | 12 | 1200 | 14400 | 0 |
|  | K59Q F159A<br>A160I |  |  |  | 1200 | 14400 | 2 |
| menitazine | WT | 3 | N/A | 3 | 200 | 600 | 0 |
|  | K59Q S92I<br>F108V Y120A<br>A160V V163F |  |  |  | 100 | 300 | 13 |
|  | K59Q S92F<br>F108V Y120A<br>A160M V163F |  |  |  | 100 | 300 | 4 |
| isotonitazene | WT | 9 | 0.5 | 3 | 200 | 600 | 0 |
|  | K59Q S92M<br>F108V Y120A<br>F159A |  |  |  | 200 | 600 | 1 |
|  | K59Q L87A<br>S92F F108V<br>Y120A A160V<br>V163F |  |  |  | 200 | 600 | 2 |

**Table S4.**
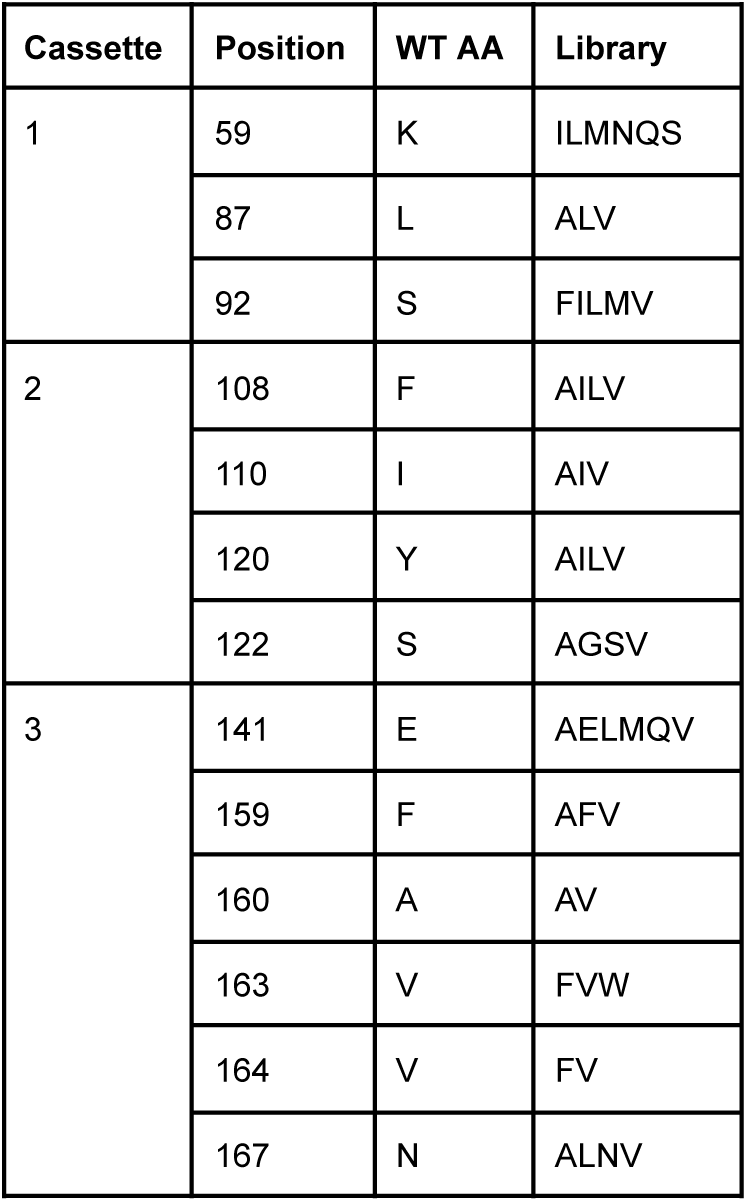
NitazeneLib1 library design. Library assembly was performed using Golden Gate of individual cassettes covering linear stretches of the PYR1 gene.

| Cassette | Position | WT AA | Library |
| --- | --- | --- | --- |
| 1 | 59 | K | ILMNQS |
|  | 87 | L | ALV |
|  | 92 | S | FILMV |
| 2 | 108 | F | AILV |
|  | 110 | I | AIV |
|  | 120 | Y | AILV |
|  | 122 | S | AGSV |
| 3 | 141 | E | AELMQV |
|  | 159 | F | AFV |
|  | 160 | A | AV |
|  | 163 | V | FVW |
|  | 164 | V | FV |
|  | 167 | N | ALNV |

**Table S5.** Theoretical coverage of NitazeneLib1 library. NitazeneLib1 was designed as three separate libraries each containing combinations of 2 out of 3 total cassettes. These sub libraries were transformed independently into *E. coli* before pooling as the final library NitazeneLib1.

|  | Library A<br>(cassettes 1&2) | Library B<br>(cassettes 1&3) | Library C<br>(cassettes 2&3) |
| --- | --- | --- | --- |
| Theoretical library size | 17,280 | 165,888 | 77,760 |
| total # *E. coli* transformants | 410,000 | 170,000 | 540,000 |
| % GFP Negative | 0.91 | 0.96 | 0.63 |
| Number of sequencing reads | 461387 | 615965 | 464819 |
| Number of sequencing reads that map to the library | 261443 | 461840 | 319432 |
| Sequences represented in library | 15673/17280<br>(90.7%) | 100422/165888<br>(60.5%) | 65416/77760<br>(84.1%) |

**Table S6.** NitazeneLib1 library input and nitazene screen output.

|  | K59 | V81 | V83 | L87 | S92 | E94 | F108 | I110 | Y120 | S122 | E141 | F159 | A160 | V163 | V164 | N167 |
| --- | --- | --- | --- | --- | --- | --- | --- | --- | --- | --- | --- | --- | --- | --- | --- | --- |
| NitazeneLib1 | ILMNQS | V | V | ALV | FILMV | E | AILV | AIV | AILV | AGSV | AELMQV | AFV | AV | FVW | FV | ALNV |
| Butonitazene M_100.1 | N |  |  | A | F |  | V |  | A |  | A |  |  | F |  | A |
| Butonitazene M_100.3 | M | L |  |  | I |  | V |  | A |  | A |  | V | F |  |  |
| Butonitazene M_100.4 | V |  |  |  | L |  | V |  | A |  | A |  | V | F |  | A |
| Butonitazene M_100.5 | V |  |  | V | I |  | V |  | A |  | A |  | V | F |  | V |
| Butonitazene M_100.6 | V |  |  |  | M |  | V |  | A |  | A | A | V | F |  | A |
| Butonitazene M_100.8 | I |  |  |  | I |  | V |  | A |  | A |  |  | F |  | A |
| Butonitazene M_100.9 | V | L |  | A | F |  | V |  | A |  | A |  | V |  |  | V |
| Butonitazene M_100.10 | N |  |  | A | F |  | V |  | A |  | A |  |  | F |  | V |
| Butonitazene M_100.12 | Q |  |  |  | M |  | I |  | L | A | A | V |  | F |  | A |
| Butonitazene M_100.14 | I |  |  |  | L |  | V |  | A |  | T |  |  | F |  | A |
| Butonitazene M_100.15 | V |  |  |  | L |  | V |  | A |  | A | A | V | F |  | A |
| Butonitazene M_100.16 | I |  |  | V | M |  | V |  | A |  | T | I | L | I |  | A |
| Isotonitazene M_100.1 | V |  |  | V | F |  | V |  | A |  | A |  | V |  |  | A |
| Menitazene M_10.1 | Q |  |  |  | M |  | L | V | A | G |  | A | V |  |  |  |
| Menitazene M_10.2 | Q |  |  |  | M |  | L | V | A | G |  | V | V | F |  |  |
| Menitazene M_100.1 | Q |  |  |  | M |  | I | V | A |  |  | V | V | F |  |  |
| Menitazene M_100.2 | Q |  |  |  | M |  | V | V | A | G |  | A | V | F |  |  |
| Menitazene M_100.5 | Q |  |  |  | M |  | V |  | A | G |  | A | V |  |  |  |
| Menitazene M_100.6 | Q |  |  |  | M |  | A |  | A | G |  |  | V | F |  |  |
| Menitazene M_100.7 | Q |  |  |  | M |  | A | V | A | G |  | V | V | F |  |  |
| Menitazene M_100.9 | Q |  |  |  | M |  | L |  | A | G |  | A | V | F |  |  |
| Menitazene M_100.10 | Q |  |  |  | M |  | V | V | A | G |  | A | V |  |  |  |
| Menitazene M_100.13 | Q |  |  |  | M |  | V |  | A | G |  | A | V | F |  |  |
| Metonitazene M_100.1 | Q |  |  |  | M |  | L | V | A | G |  | A | V |  |  |  |
| Metonitazene M_100.3 | Q |  |  |  | M |  | L | V | A | G |  | V | V | F |  |  |
| Metonitazene M_100.4 | Q |  |  |  | M |  | I | V | A | G |  | A | V | F |  |  |
| Metonitazene M_100.5 | Q |  |  |  | M |  | V | V | A | G |  | A | V |  |  |  |
| Metonitazene M_100.7 | Q |  |  |  | M |  | L |  | A | G |  | A | V | F |  |  |
| Nitazene M_N1 | Q |  |  |  | M |  | V |  | A | G |  | A | V |  |  |  |
| Nitazene M_N7 | Q |  |  |  | M |  | V |  | A | A |  | A | V |  |  |  |
| Nitazene M_N8 | Q |  |  |  | M |  | V |  | A |  |  | A | V |  |  |  |
| Nitazene M_N13 | Q |  |  |  | M |  | V | V | A | G |  | A | V |  |  |  |
| Nitazene M_N14 | Q |  |  |  | M |  | V |  | A | G |  | V | V | F |  |  |
| Nitazene M_10.2 | Q |  |  |  | M |  | L | V | A | G |  | A | V |  |  |  |
| Nitazene M_100.1 | Q |  |  |  | M |  | V |  | P |  |  | V | V | F |  |  |
| Nitazene M_100.5 | Q |  |  |  | M |  | L | V | A | G |  | V | V | F |  |  |
| Nitazene M_100.6 | Q |  |  |  | M |  | V |  | A | G |  |  | V | F | F |  |
| Nitazene M_100.9 | I |  |  |  | M |  | V |  | A |  |  | A | V |  |  |  |
| Nitazene M_100.10 | Q |  |  |  | M |  | L |  | A | G |  | A | V | F |  |  |
| Nitazene M_100.11 | M |  |  |  | F |  | V |  | A |  |  |  | V | F |  |  |
| Protonitazene M_100.1 | V | L |  |  | A | I | V |  | A |  | Q | V | V | F |  | V |
| Protonitazene M_100.4 | V |  |  |  | M |  | V |  | A |  | A | A | V | F |  | V |
| Protonitazene M_100.5 | Q |  |  |  | M |  | V |  | A | G |  | A | V | F |  |  |
| Protonitazene M_100.6 | V |  |  | V | F |  | V |  | A |  | V |  | V |  |  | V |
| Protonitazene M_100.7 | I |  |  |  | I |  | V |  | A |  | A |  |  | F |  | A |
| Protonitazene M_100.8 | V |  |  |  | L |  | V |  | A |  | A | A | V | F |  | A |
| Protonitazene M_100.9 | V |  |  |  | M |  | V |  | A |  | A | A | V | F |  | A |
| Protonitazene M_100.10 | I | L |  |  | M |  | V |  | A |  | A | A | V | F |  | A |
| Protonitazene M_100.11 | Q |  |  |  | M |  | V |  | A | G |  | A | V |  |  |  |

**Table S7.** Nitazene Library 2 mutations.

| Ultramer | Position | PYR1<sup>nita</sup> AA | Library AA | Codon |
| --- | --- | --- | --- | --- |
| Forward | 81 | V | VIL | VTT |
|  | 83 | V | VILM | VTK |
|  | 92 | M | MILF | WTK |
|  | 108 | V | VI | RTH |
| Reverse 1 | 120 | A | GA | GSS |
|  | 122 | G | DEAG | GVN |
|  | 141 | E | DE | GAW |
|  | 159 | A | GA | GYG |
| Reverse 2 | 159 | A | H | CAT |

**Table S8.** Nitazene Library 3 and 4 mutations.

| Ultramer | Position | PYR1<sup>nita2</sup> AA | PYR1<sup>nita2.1</sup> AA | Nitazene Library 3 AA | Nitazene Library 4 AA | Codon |
| --- | --- | --- | --- | --- | --- | --- |
| Forward | 59 | Q | Q | QVASLIMT | QVASLIMT | DYR |
|  | 83 | L | V | V | L |  |
|  | 92 | M | F | F | M |  |
|  | 120 | A | G | G | A |  |
| Reverse 1 | 122 | G | E | E | G |  |
|  | 141 | D | D | EDAG | EDAG | GVN |
|  | 159 | H | H | VA | VA | GYT |
|  | 163 | V | V | VA | VA | GYN |
|  | 164 | V | V | VILM | VILM | VTK |
|  | 167 | N | N | SACGN | SACGN | KSY |
| Reverse 2 | 159 | H | H | H | H |  |

## Notes

doi:10.5281/zenodo.15298585

